# Cytoplasmic redox imbalance in the thioredoxin system activates Hsf1 and results in hyperaccumulation of the sequestrase Hsp42 with misfolded proteins

**DOI:** 10.1101/2023.06.26.546610

**Authors:** Davi Gonçalves, Sara Peffer, Kevin A. Morano

## Abstract

Cells employ multiple systems to maintain homeostasis when experiencing environmental stress. For example, the folding of nascent polypeptides is exquisitely sensitive to proteotoxic stressors including heat, pH and oxidative stress, and is safeguarded by a network of protein chaperones that concentrate potentially toxic misfolded proteins into transient assemblies to promote folding or degradation. The redox environment itself is buffered by both cytosolic and organellar thioredoxin and glutathione pathways. How these systems are linked is poorly understood. Here, we determine that specific disruption of the cytosolic thioredoxin system resulted in constitutive activation of the heat shock response in *Saccharomyces cerevisiae* and accumulation of the sequestrase Hsp42 into an exaggerated and persistent juxtanuclear quality control (JUNQ) compartment. Terminally misfolded proteins also accumulated in this compartment in thioredoxin reductase (*TRR1*)-deficient cells, despite apparently normal formation and dissolution of transient cytoplasmic quality control (CytoQ) bodies during heat shock. Notably, cells lacking *TRR1* and *HSP42* exhibited severe synthetic slow growth exacerbated by oxidative stress, signifying a critical role for Hsp42 under redox challenged conditions. Finally, we demonstrated that Hsp42 localization patterns in *trr1Δ* cells mimic those observed in chronically aging and glucose-starved cells, linking nutrient depletion and redox imbalance with management of misfolded proteins via a mechanism of long-term sequestration.

## Introduction

Cells respond to fluctuating external and internal environmental changes through physiological adaptation driven in part by transcriptional changes in gene expression. When the stressors are proteotoxic and impact the folding, maturation or function of the proteome, an array of cytoprotective genes that includes protein molecular chaperones, components of the ubiquitin-proteasome system and detoxification enzymes are produced in part through the action of the heat shock transcription factor, Hsf1 (Akerfelt *et al*., 2010). In the yeast *Saccharomyces cerevisiae*, such stressors include heat shock, amino acid analogs, glucose starvation and oxidative stress (Trotter *et al*., 2002; Verghese *et al*., 2012; Tye and Churchman, 2021). In addition to oxidizing agents like hydrogen peroxide and superoxide anions that damage proteins, a range of thiol-reactive compounds are potent inducers of Hsf1 and the resultant heat shock response (HSR) (West *et al*., 2012). Multiple chemical mechanisms of thiol-reactive proteotoxic stress have been documented, including inappropriate disulfide bond formation, irreversible oxidation, and adduct formation on sulfur atoms within the amino acids methionine and cysteine (Dahl *et al*., 2015; Fra *et al*., 2017; Lévy *et al*., 2019). The relative pKa of the cysteine thiolate anion is determined by the local microenvironment and defines the reactivity of the side chain to thiol-reactive stressors; most cysteines are non-reactive while some protein cysteines are hypersensitive to insult (Winterbourn and Hampton, 2008). For example, the heat shock protein (Hsp) 70 family of chaperones possesses multiple highly reactive cysteines whose modification inhibits chaperone function (Miyata *et al*., 2012; Wang *et al*., 2012; Wang and Sevier, 2016). Moreover, in yeast, the Hsp70 Ssa1 governs the response of Hsf1 to thiol-reactive stress via two cysteine triggers (C264, C303) in the ATPase domain that result in release of the chaperone from Hsf1 intrinsically disordered domains required for three-dimensional chromatin organization and transcriptional activation (Wang *et al*., 2012; Santiago and Morano, 2022).

The linked but independent thioredoxin and glutathione systems control the redox state of the yeast cytoplasm. Electrons produced from catabolic oxidation (i.e., glycolysis, the tricarboxylic acid cycle and the pentose phosphate pathway) are shuttled through thioredoxin (Trr1) or glutathione (Glr1) reductases to thioredoxin (Trx1/2) or glutathione (GSH), respectively, that partner with peroxiredoxins to collaboratively maintain the cytoplasm in a reduced state distinct from the oxidizing environment of the endomembrane system (endoplasmic reticulum, Golgi, lysosome/vacuole) and the extracellular space (Le Moan *et al*., 2006; López-Mirabal and Winther, 2008; Ayer *et al*., 2014). The glutathione system has been defined as the first line of antioxidant defense in budding yeast, which maintain high concentrations of this small molecule to detoxify thiol reactive agents (Spector *et al*., 2001; Le Moan *et al*., 2006). In contrast, the thioredoxin system plays the dominant role in maintaining cellular redox homeostasis (Trotter and Grant, 2003).

The proteome is surveilled in real time via the action of the protein quality control network, comprised primarily by chaperones that recognize inappropriately exposed hydrophobic surfaces of unassembled complex subunits or regions of misfolded proteins. Once identified, these substrates are collected into transient assemblies known as Q-, or CytoQ-bodies, which include Hsp70 chaperones (Ssa1/2/3/4) as well as Hsp40 (Sis1, Ydj1) and Hsp110 (Sse1/2) co-chaperones, the disaggregase Hsp104, and other components including small heat shock proteins (sHSP) (Kaganovich *et al*., 2008; Malinovska *et al*., 2012; Miller *et al*., 2015b; Sathyanarayanan *et al*., 2020). It has been recently established that at least two sHSPs, Btn2 and Hsp42, function as “sequestrases” that intercalate between misfolded chaperone cargo to generate the CytoQ assemblies by virtue of intrinsically disordered/prion-like domains (Malinovska *et al*., 2012; Miller *et al*., 2015a; Grousl *et al*., 2018; Shrivastava *et al*., 2022). CytoQ assemblies fail to form in cells lacking Hsp42, underscoring the important role the sequestrase plays in shepherding misfolded proteins (Specht *et al*., 2011; Saarikangas and Barral, 2015). Multiple CytoQ bodies typically form upon proteotoxic stress, which then resolve into one or more possible depots: the internuclear quality control compartment (INQ), the cytoplasmic, juxtanuclear quality control compartment (JUNQ), or in the case of amyloid deposits and prion-like proteins, the insoluble protein deposit (IPOD) (Kaganovich *et al*., 2008; Miller *et al*., 2015b; Hill *et al*., 2017; Rothe *et al*., 2018). INQ/JUNQ are marked by substrates that are subsequently extracted by Hsp104 and either refolded or ubiquitinated as a prelude to degradation (Specht *et al*., 2011; Mathew *et al*., 2017). The Btn2 sequestrase localizes to the nucleoplasm and is required for INQ formation, while Hsp42 appears to be restricted to the cytoplasm and promotes formation of JUNQ and IPOD (Malinovska *et al*., 2012; Miller *et al*., 2015a, 2015b). INQ and JUNQ only accumulate when protein degradation is blocked and do so as one or two large static foci in cells where model misfolded proteins fused to GFP are tracked via fluorescence microscopy (Kaganovich *et al*., 2008; Kumar *et al*., 2022).

In this report, we use a genetic approach to generate redox stress in yeast cells via targeted deletion of major gene products comprising the thioredoxin and glutathione pathways. Strikingly, we discovered that proteotoxic stress as reported by Hsf1 activation is only experienced by cells with a disrupted cytosolic, but not mitochondrial, thioredoxin system, with no apparent role for the glutathione pathway. We found that the sequestrase Hsp42, but not Btn2 or the other yeast sHSP Hsp26, hyperaccumulated in 1-2 large foci in *trr1Δ* cells in a structure co-localizing with the nuclear membrane, consistent with JUNQ compartments. The foci formed in *trr1Δ* cells were independent of heat shock-induced transient CytoQ bodies and did not include the Hsp104 or Ssa/Ssb Hsp70 chaperones. While protein folding and degradation were not impaired in *trr1Δ* cells as reported using model substrates, permanently misfolded proteins colocalized with Hsp42, further evidence of aberrant quality control in cells lacking thioredoxin reductase function. Importantly, Hsp42 sequestration was shown to be important for optimal growth and survival in unstressed or oxidant-challenged *trr1Δ* cells. Finally, Hsp42 dynamics in *trr1Δ* cells were found to closely mirror localization patterns in glucose-starved and aging cells, suggesting a common underlying mechanism that connects redox balance, abundance of reducing equivalents, and spatial quality control.

## Results

### Cells deficient in the thioredoxin system exhibit constitutive activation of the HSR

Based on previous work demonstrating that thiol-reactive compounds are potent activators of the HSR, we sought a genetic approach to alter redox balance in the absence of exogenous insults (Trott *et al*., 2008; Wang *et al*., 2012). To accomplish this, cells lacking the *TRR1* and/or redundant *TRX1* and *TRX2* genes were transformed with a β-galactosidase transcriptional reporter plasmid driven by a typical heat shock element containing minimal promoter (HSE-*lacZ*). This reporter specifically monitors Hsf1 transcriptional activity and has been successfully used as a sensitive indicator of HSR status (Bonner *et al*., 1994; Morano *et al*., 1999). Reporter activity was strikingly elevated in *trr1Δ*, *trx1Δtrx2Δ*, and *trr1Δtrx1Δtrx2Δ* cells as compared to wild type cells under non stress conditions (Fig. 1A). While the *trr1Δ* and *trx1Δtrx2Δ* strains exhibited nearly identical elevated levels of constitutive HSE-*lacZ* activity, the triple mutant displayed a lower, but still heightened, level of activation. To further explore the connections between cellular redox buffering systems and HSR regulation, we assessed HSE-*lacZ* in additional genetic backgrounds. Consistent with their redundant nature, cells lacking either the thioredoxin genes *TRX1* or *TRX2* exhibited no change in the HSR (Fig. S1). A similar outcome was observed in cells deficient in the mitochondrial thioredoxin reductase system (*trr2Δ* and *trx3Δ*) (Trotter and Grant, 2005). The methionine sulfoxide reductase Mxr1 is required to reduce oxidized methionine residues (Kaya *et al*., 2010). Deletion of *MXR1* likewise did not affect HSE-*lacZ* activity. We additionally tested the role of the glutathione system in HSR regulation. Loss of the glutathione reductase *GLR1* did not activate the HSR. Synthesis of glutathione requires the gene *GSH1*, encoding gamma glutamylcysteine synthetase, and *gsh1Δ* mutants are inviable in the absence of exogenous reduced glutathione (Spector *et al*., 2001). We therefore cultured *gsh1Δ* cells in the presence of 1 mM glutathione, followed by washout and growth for an additional six hours in the presence or absence of glutathione and observed no change in HSE-*lacZ* activity. We confirmed the reporter assay results by measuring endogenous transcript levels of the *SSA3* and *SSA4* Hsp70 genes via qRT-PCR (Fig. 1B). Together, these data demonstrate that the cytoplasmic thioredoxin system is uniquely linked to Hsf1 transcriptional activation in *Saccharomyces cerevisiae* cells grown under optimal, normoxic conditions.

**Figure 1.**
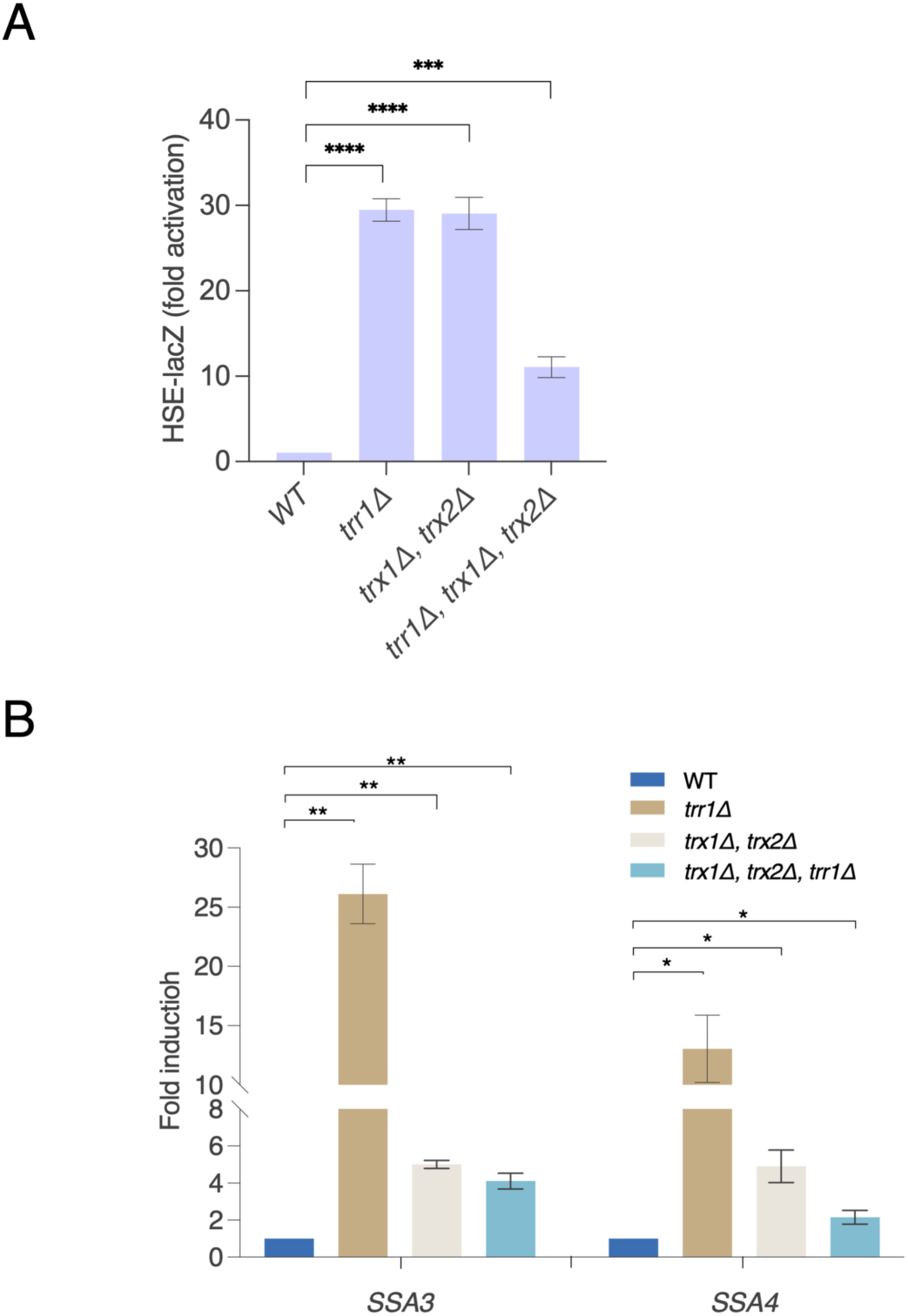
Cells deficient in the thioredoxin system exhibit constitutive activation of the HSR. A) The indicated strains bearing the pSSA3HSE-lacZ reporter plasmid were grown to mid-log phase and β-galactosidase activity measured as described in the Materials and Methods. B) The indicated strains were grown to mid-log phase, RNA extracted and levels of the *SSA3* and *SSA4* transcripts determined by qRT-PCR using the *TAF10* gene as a normalization control. Statistical significance between the indicated strains was determined using Welch’s unpaired t test (p=0.05, *; p=0.005, **; p=0.0005, ***; p=0.00005, ****).

### Thioredoxin system mutants hyperaccumulate the sequestrase Hsp42

The observed chronic HSR activation in thioredoxin-mutant cells suggested that one or more aspects of general cellular proteostasis is negatively impacted in the absence of the thioredoxin system. Previous work has established that misfolded proteins are collected into transient deposition sites for storage and/or processing, which may include ubiquitination and eventual degradation by the proteasome (Miller *et al*., 2015b). One of the first steps in this process for cytoplasmic proteins is the formation of Q-bodies (also called CytoQ) that typically include the sequestrase Hsp42, one or more Hsp70 proteins (Ssa1, Ssa2, Ssa3, Ssa4) and their cofactors (Sse1/2, Ydj1, Sis1), and the disaggregase Hsp104. Q-bodies containing misfolded proteins either resolve via refolding catalyzed by the abovementioned chaperones or coalesce into larger depots termed juxtanuclear (JUNQ) or intranuclear (INQ) quality control compartments (Kaganovich *et al*., 2008; Miller *et al*., 2015b; Hill *et al*., 2017). We employed fluorescence microscopy to assess the status of multiple chaperone proteins using chromosomally encoded green fluorescent protein (GFP) fusions. Strikingly, while diffuse in wild type cells, Hsp42-GFP accumulated at a high frequency (40-60% of cells) in large, generally single foci in cells lacking cytoplasmic thioredoxin reductase, both cytoplasmic thioredoxins, or all three proteins (Fig. 2A). Cells expressing GFP alone did not exhibit concentration of fluorescence signal. Because we observed no significant differences between the *trr1Δ* and *trx1Δtrx2Δ* strain backgrounds in our studies, all further experiments were carried out using the former deletion strain to inactivate the thioredoxin system. GFP fusions to the Hsp70s Ssa1, Ssa4, Ssb1 and the Hsp40 chaperone Ydj1 also remained diffuse in *trr1Δ* cells (Fig. 2B). The chaperones Btn2 and Hsp26, as members of the sHSP family, contain both conserved α-crystallin chaperone domains as well as divergent amino- and carboxyl terminal domains that dictate differential behavior with respect to substrate recruitment (Fig. S2A) (Mogk *et al*., 2019). Additionally, Btn2 possesses protein sequestrase activity similar to Hsp42 (Malinovska *et al*., 2012; Miller *et al*., 2015a). However, GFP fusions to both proteins remained diffusely localized in *trr1Δ* cells (Fig. 2B). The cellular levels of Btn2 and Hsp26 are quite low in unstressed wild type cells, but significantly increased in *trr1Δ* cells, as are levels of Hsp42 as assessed by immunoblot (Fig. S2B). These results are consistent with all three being under the transcriptional control of Hsf1 and suggest that the presence of the large Hsp42 foci is not due to overexpression of the chaperone (Solís *et al*., 2016). To confirm that the altered localization pattern of Hsp42 was due to the enzymatic activity of Trr1 and its role in maintaining proper redox balance in the cytoplasm, we generated an Hsp42-GFP strain wherein the catalytic and resolving cysteine residues of Trr1 required for the disulfide exchange reaction were substituted with serine (Chae *et al*., 1994). Cells expressing the *trr1-C142S, C145S* allele exhibited the same high level of Hsp42-GFP focus formation as *trr1Δ* cells bearing an empty vector, as well as the drastic slow-growth phenotype of cells lacking thioredoxin reductase activity (Fig. S3A, B) (Trotter *et al*., 2002; Kritsiligkou *et al*., 2018).

**Figure 2.**
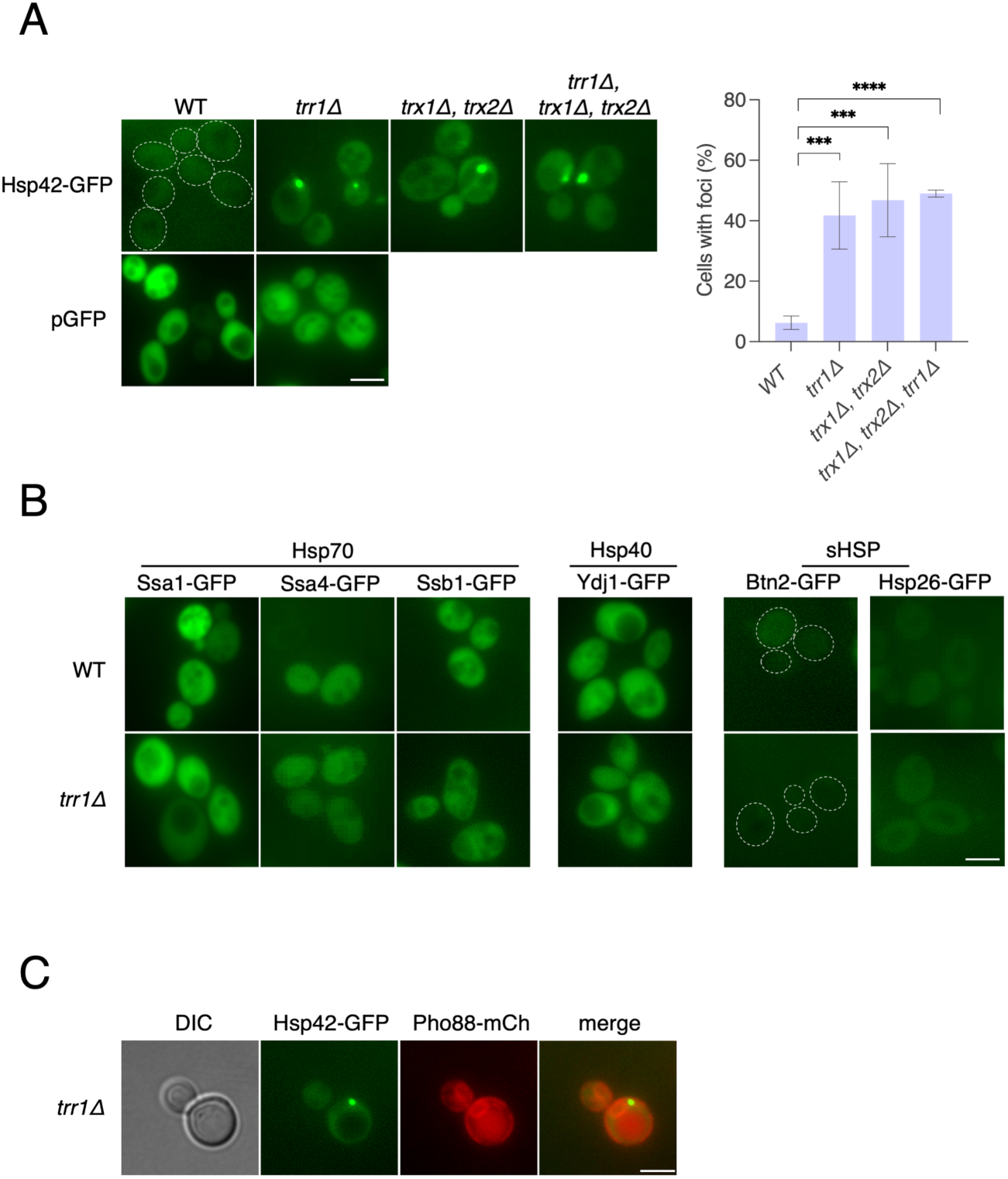
Thioredoxin system mutants hyperaccumulate the sequestrase Hsp42. A) The indicated strains with GFP integrated in-frame with *HSP42*, or transformed with a plasmid expressing GFP alone, were grown to mid-log phase and imaged as described in Materials and Methods. The percentage of cells with foci was determined by counting at least 100 cells from multiple fields. B) Wild type (WT) or *trr1Δ* cells expressing GFP fusions to the indicated genes at the genomic locus were imaged in mid-log phase. C) Strain HSP42-GFP *trr1Δ* PHO88-mCh was grown to mid-log phase and imaged as described in Materials and Methods. Statistical significance between the indicated strains was determined using Welch’s unpaired t test (p=0.05, *; p=0.005, **; p=0.0005, ***; p=0.00005, ****). Scale bar = 5 µm.

In addition to the JUNQ and INQ compartments, a third depot termed the IPOD has been described in yeast that typically includes amyloid-forming proteins as well as Hsp42 and other chaperones (Kaganovich *et al*., 2008; Rothe *et al*., 2018). While JUNQ and INQ are perinuclear and intranuclear, respectively, IPOD is typically peri-vacuolar or randomly localized within the cytoplasm. To assess where Hsp42-GFP was localizing in *trr1Δ* cells, we constructed a strain expressing an mCherry-tagged allele of the ER membrane protein Pho88 that also outlines the nuclear membrane (D’Urso *et al*., 2016). The Hsp42-GFP signal was observed to be immediately adjacent to the cytoplasmic face of the nuclear membrane as judged by comparison to the Pho88-mCh signal, consistent with accumulation/persistence of a JUNQ-like compartment in *trr1Δ* cells (Fig. 2C).

### Chaperone and substrate dynamics in trr1Δ cells

To further explore the nature of the large Hsp42 foci in *trr1Δ* cells, we assessed chaperone localization in response to protein misfolding stress. Wild type and *trr1Δ* cells bearing either Hsp42-GFP or a red fluorescent protein (RFP) fusion to the disaggregase Hsp104 (Hsp104-RFP) were grown at the non-stress temperature of 30°C and shifted or not to 39°C (HS) for 20 min. Heat-shocked cells were then returned to 30°C for a five hour recovery period. As expected, wild type cells exhibited diffuse fluorescence in control conditions, and formed multiple small foci in nearly all cells (CytoQ) during heat shock (Fig. 3A). These CytoQ bodies were largely resolved into a single focus in approximately 40% of cells during the recovery period. In *trr1Δ* cells, the same CytoQ dynamics were observed, while the single large foci remained present during all phases of the experiment. Interestingly, the persistence of large, single foci, presumably exaggerated JUNQ compartments, was greater in *trr1Δ* cells than in wild type cells (over 80% vs. ∼ 40%) at the end of the recovery period, suggesting that the resolved CytoQ bodies were “added” to the redox-dependent body. This possibility is further supported by the fact that we did not observe two independent large foci in Hsp42-GFP *trr1Δ* cells after the recovery phase. We performed the same experiment using Hsp104-RFP-containing cells and noted that while both wild type and *trr1Δ* strains accumulated CytoQ during the heat shock, in neither strain were persistent large foci observed under control or recovery conditions (Fig. 3B). These findings establish that Hsp42, like Hsp104, is rapidly recruited to CytoQ structures under protein misfolding stress; however, Hsp42 fails to disengage during the recovery phase in contrast to Hsp104. Moreover, in *TRR1*-deficient cells, CytoQ dynamics proceed normally with the addition of persistent JUNQ compartments containing Hsp42.

**Figure 3.**
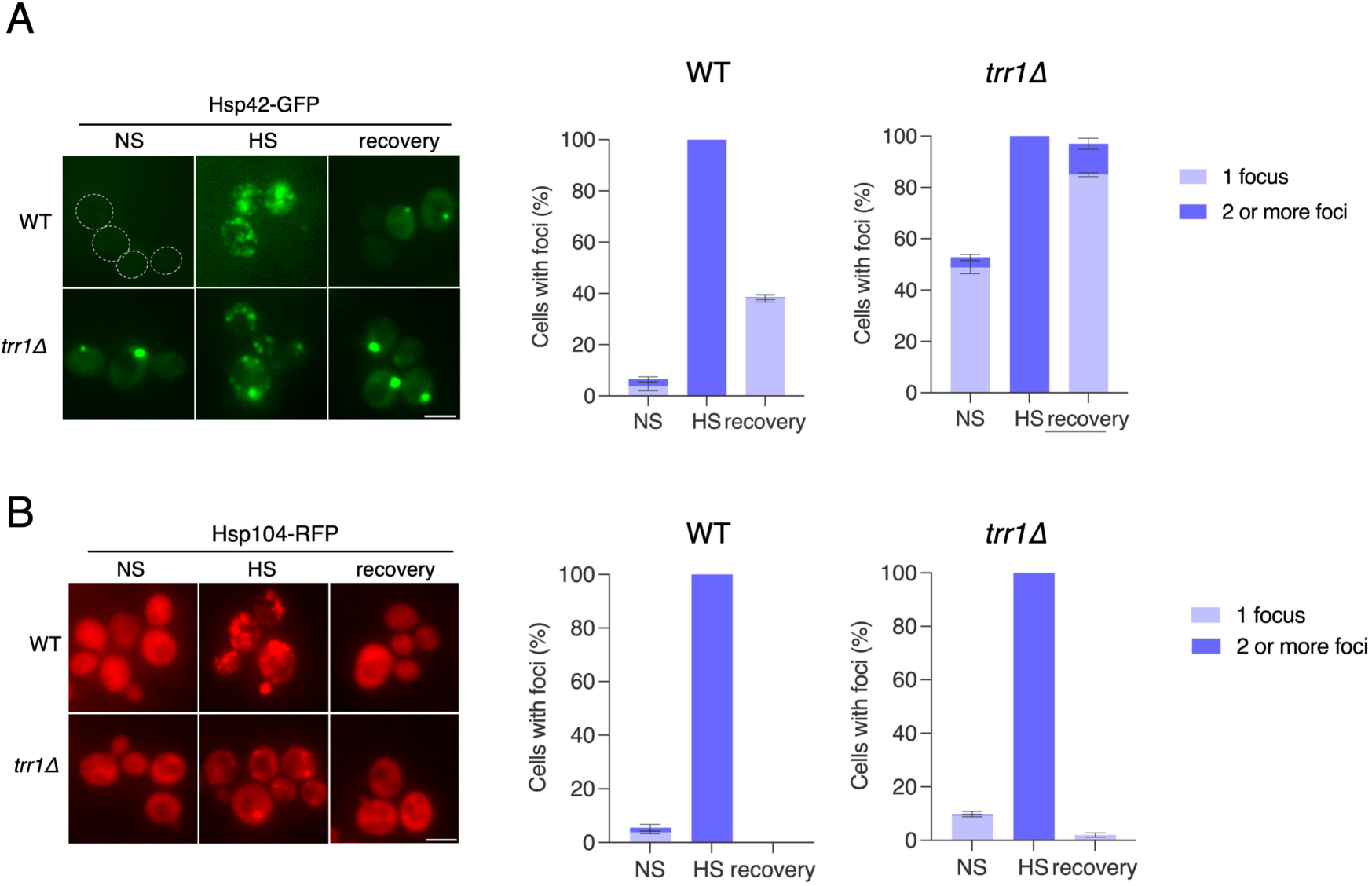
Chaperone and substrate dynamics in *trr1Δ* cells. The indicated strains expressing Hsp42-GFP (A) or Hsp104-RFP (B) from the chromosomal locus were grown to mid-log phase at 30°C (NS), shifted to 39°C for 20 min (HS), and allowed to recover for 5 h at 30°C (recovery). The percentage of cells with one or two or more foci was determined by counting at least 100 cells from multiple fields. Scale bar = 5 µm.

### Terminally misfolded proteins accumulate in Hsp42-GFP foci in trr1Δ cells

The Hsp42 sequestrase forms multimeric scaffolded structures with misfolded proteins that both prevents their aggregation and facilitates extraction and refolding as well as degradation (Mogk and Bukau, 2017). The persistence of JUNQ compartments containing Hsp42-GFP in cells defective in the thioredoxin system suggested that misfolded proteins might also accumulate within these structures. To test this hypothesis, we expressed the terminally misfolded model protein CPY^‡^-GFP in wild type and *trr1Δ* cells and monitored localization dynamics in the presence or absence of the translation inhibitor cycloheximide (Heck *et al*., 2010). CPY^‡^-GFP typically forms small CytoQ bodies that are resolved over time via degradation by the proteasome (Fig. 4A). In contrast, in *trr1Δ* cells, CPY^‡^-GFP accumulated in 1-2 large foci, and, after 90 min of cycloheximide chase, the majority of cells exhibited a single very large focus reminiscent of the Hsp42-GFP/JUNQ compartment. These results suggest that the CPY^‡^-GFP within the CytoQ bodies failed to be extracted and degraded in *trr1Δ* cells and instead hyperaccumulated into an exaggerated JUNQ compartment. Indeed, when the same experiment was performed using the proteasome inhibitor MG-132 instead of cycloheximide, both wild type and *trr1Δ* cells displayed similar accumulation of two or more exaggerated foci, consistent with a degradation block (Fig. 4B). Accumulation of CPY^‡^-GFP into foci of any size was entirely dependent on Hsp42 as previously reported, as no foci were detected in both *TRR1 hsp42Δ* and *trr1Δ hsp42Δ* cells (Fig. 4C) (Specht *et al*., 2011; Saarikangas and Barral, 2015). Finally, to verify that the Hsp42-GFP and misfolded protein foci accumulating in *trr1Δ* cells are one and the same, we generated an Hsp42-RFP fusion protein and co-expressed it with either CPY^‡^-GFP or another permanently misfolded substrate, the truncated Gnd1 protein (tGND-GFP) (Heck *et al*., 2010). In both cases, the foci completely overlapped. Together, these results demonstrate that the JUNQ compartments persisting in *trr1Δ* cells contain both the sequestrase Hsp42 as well as misfolded proteins. These proteins are likely associated with each other in a manner that precludes degradation in the case of the two model proteins and refolding in the case of endogenous substrates normally processed through transient CytoQ assemblies.

**Figure 4.**
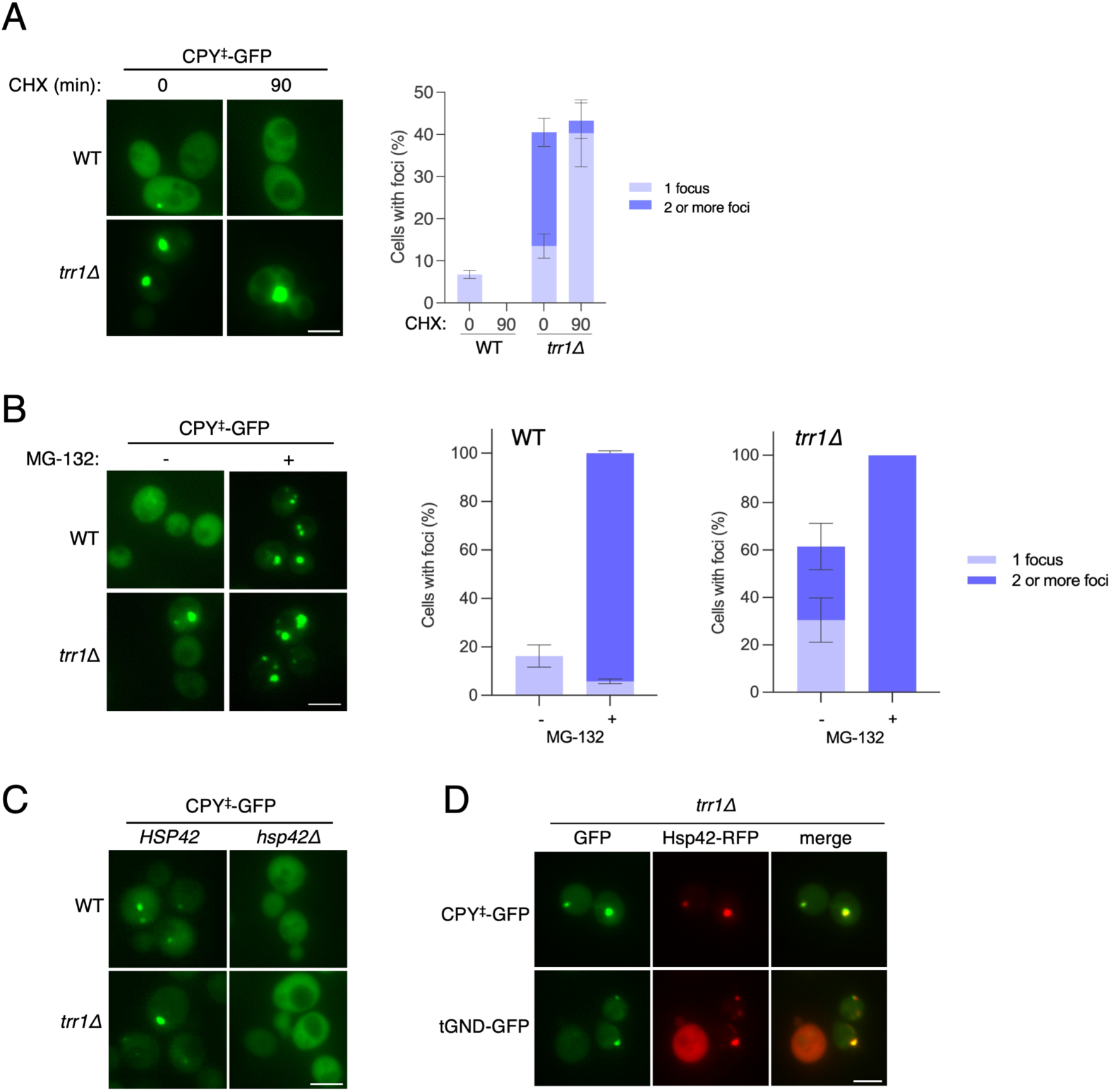
Terminally misfolded proteins accumulate in Hsp42-GFP foci in *trr1Δ* cells. A) The indicated strains expressing the CPY^‡^-GFP misfolded protein fusion from the chromosome were grown at 30°C and imaged immediately prior to (0) or 90 min after (90) treatment with cycloheximide as described in Materials and Methods. The percentage of cells with one or two or more foci was determined by counting at least 100 cells from multiple fields. B) The indicated strains (additionally *pdr5Δ*) were grown at 30°C in the absence (-) or presence (+) of MG-132 for 2 h. The percentage of cells with one or two or more foci was determined by counting at least 100 cells from multiple fields. C) The indicated strains were grown to mid-log phase at 30°C and imaged. D) Strains expressing Hsp42-RFP as well as either CPY^‡^-GFP or tGND-GFP were grown to mid-log phase and imaged in both the red and green channels. Scale bar = 5 µm.

One explanation for the persistent JUNQ structures could be excess production of misfolded proteins in *trr1Δ* cells; i.e., an overall increase in protein misfolding due to an altered redox environment. To test this possibility, we expressed the thermally labile protein firefly luciferase (FFL) as a GFP fusion protein in wild type and *trr1Δ* cells (Abrams and Morano, 2013). FFL-GFP misfolds under proteotoxic stress conditions and localizes to transient CytoQ bodies (Tkach and Glover, 2008; Escusa-Toret *et al*., 2013; Miller *et al*., 2015a). We therefore heat-shocked FFL-GFP-bearing cells, followed by return to normal temperature in the presence of cycloheximide, and assessed both luciferase enzymatic activity and focus formation. FFL-GFP recovered approximately 50% of initial activity in wild type cells over 90 min, with a moderately faster and higher rate of recovery in *trr1Δ* cells (Fig. S4A). Consistent with these results, CytoQ punctae formed in both strains during heat shock and partially resolved by 90 min to a similar degree as assessed by fluorescence microscopy (Fig. S4B). While not an exhaustive analysis, these data suggest that global protein refolding is not grossly impaired in *trr1Δ* cells.

An alternative explanation for the persistence of both misfolded protein like CPY^‡^-GFP and tGND-GFP in Hsp42-containing JUNQ compartments might be failure to properly target these substrates for degradation through the ubiquitin-proteasome system (UPS) (Baker and Bernardini, 2021). We assessed this possibility in two different ways. The addition of polyubiquitin chains to cellular misfolded proteins can be detected by immunoblot using anti-ubiquitin antibodies if downstream proteasome degradation is blocked. We therefore treated wild type and *trr1Δ* cells with either MG-132, cycloheximide or both compounds and prepared whole cell extracts for immunoblot. A characteristic smear of ubiquitin-positive proteins was observed in untreated cells whose intensity decreased with cycloheximide (due to fewer nascent chains being produced) and enhanced with both cycloheximide and MG-132 (due to proteasome inhibition) in both wild type and *trr1Δ* cells, suggesting that the ubiquitination phase of protein quality control was unaffected by redox imbalance (Fig. S5A). To ask whether proteasome activity itself might be abrogated in *trr1Δ* cells, leading to accumulation of misfolded proteins, we prepared native cell extracts and utilized a commercial proteasome enzymatic activity assay. 20S chymotrypsin-like proteasome activity was measured and validated by comparison with measurements taken in the presence of MG-132. Surprisingly, 20S proteasome activity was approximately three-fold higher in *trr1Δ* cells as compared to wild type, possibly due to enhanced proteasome gene expression driven by the *HSF1-RPN4* circuit (Fig. S5B) (Hahn *et al*., 2006). Together, these results demonstrate that overall function of the UPS is intact, if not elevated, in cells lacking thioredoxin reductase activity.

### Cells deficient in both thioredoxin reductase activity and sequestrase function experience synthetic slow growth and hypersensitivity to oxidative stress

Our experiments demonstrated that *trr1Δ* cells accumulated both Hsp42 and misfolded proteins in an exaggerated JUNQ compartment that failed to resolve despite an intact UPS. Notwithstanding a near-absolute requirement for Hsp42 to form CytoQ or JUNQ structures in yeast cells, *hsp42Δ* cells exhibit essentially no phenotypes unless cells are severely compromised for protein quality control through the Hsp70 chaperone network (Ho *et al*., 2019). However, because our findings suggested that Hsp42 may be a critical chaperone in cells that lack the ability to maintain a reduced cytoplasm, we generated an *hsp42Δ trr1Δ* strain and compared its growth to wild type and single-mutant strains. As previously reported, *trr1Δ* cells exhibited a slow growth phenotype under normal conditions, while *hsp42Δ* cells grew indistinguishably from wild type. In contrast, *hsp42Δ trr1Δ* cells exhibited profoundly slow growth (Fig. 5A). To ask whether exogenous oxidative stress may exacerbate these effects, the same four strains were cultured in the presence of 0.5 mM H_2_O_2_. A similar pattern was observed, with more intense growth retardation of both *trr1Δ* and *hsp42Δ trr1Δ* cells, the latter of which were nearly inviable. Together, these data highlight the interconnectedness of sequestrase and thioredoxin reductase functions and to our knowledge represent a rare example of synthetic growth defects resulting from genetic disruption of the *HSP42* locus in yeast.

**Figure 5.**
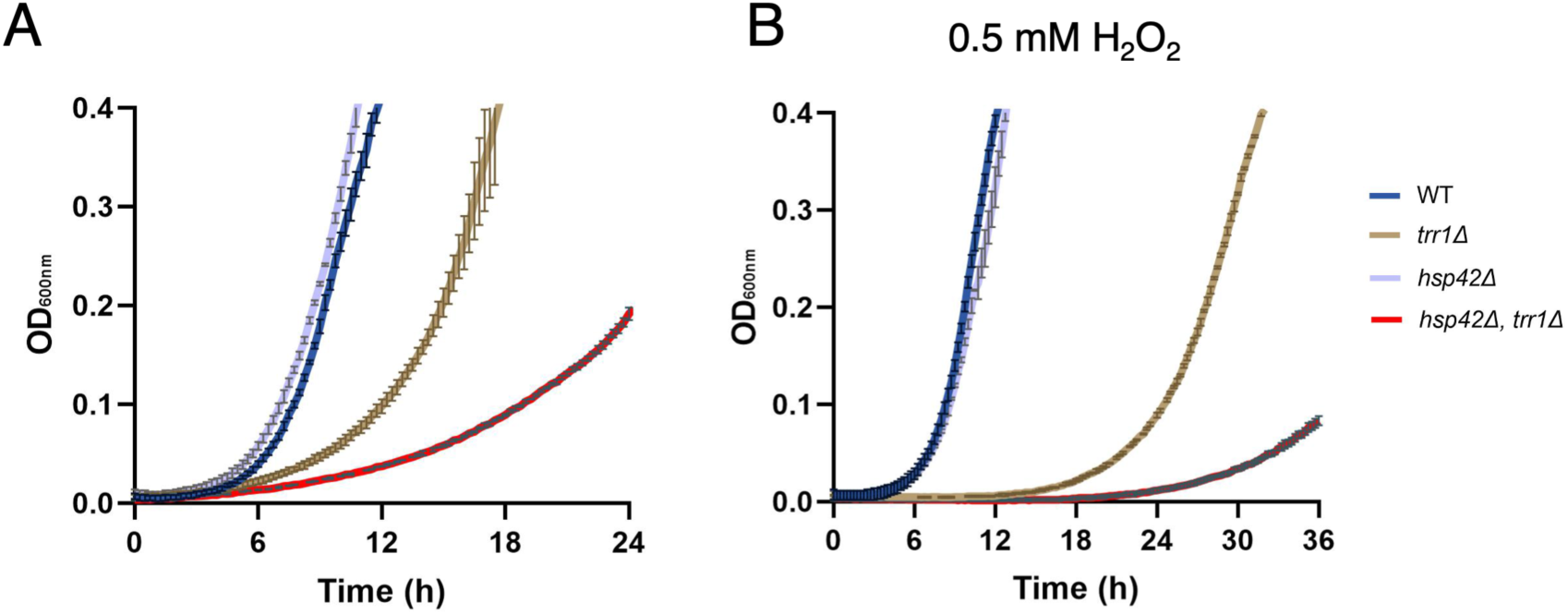
Cells deficient in both thioredoxin reductase activity and sequestrase function experience synthetic slow growth and hypersensitivity to oxidative stress. A) The indicated strains were inoculated at an initial OD_600_=0.01 in a sterile 96-well plate and grown with shaking at 30°C for 24 h with density measurements taken every 10 min. The average of three biological replicate growth curves are shown with standard deviation. B) Same as (A), but cultures were additionally grown in the presence of 0.5 mM H_2_O_2_ for 36 hr.

### Hsp42 dynamics in glucose-starved and chronologically aging cells phenocopy trr1Δ cells

Yeast growth in a fixed culture over time has been interpreted as a microbial model of eukaryotic aging; cells initially exhibit robust growth in a nutrient-rich environment that slows as key nutrients, including glucose, are exhausted (Herman, 2002). The same metabolic state can be achieved by shifting logarithmically growing, glucose-rich (2%) cultures to a minimal glucose concentration (0.02%), bringing about acute glucose starvation. Hsp42 has been previously shown to accumulate in so-called “Hsp42 bodies” in yeast cultures during stationary phase or in response to glucose starvation that bear a strong resemblance to the enhanced JUNQ structures observed in our experiments (Narayanaswamy *et al*., 2009; Liu *et al*., 2012; O’Connell *et al*., 2014; Lee *et al*., 2016). We therefore sought to better understand possible similarities between all these observations. Wild type and *trr1Δ* cells were grown in rich glucose media and then shifted to low glucose conditions and Hsp42-GFP focus accumulation assessed. While *trr1Δ* cells exhibited a high percentage of single-focus formation in both growth environments, wild type cells only accumulated an Hsp42-GFP focus under starvation conditions (Fig. 6A). In fact, although the percentage of single foci increased in the *trr1Δ* strain after shift to low glucose (50% to approximately 75%), the total percentage was nearly identical in both strains. We interpret this result to mean that *trr1Δ* cells largely phenocopy the glucose starved scenario in wild type cells and that acute starvation is minimally additive with genetically induced redox imbalance. To ask whether the same concept held true for batch culture aging, wild type and *trr1Δ* cells were grown in parallel cultures and images taken of Hsp42-GFP localization at defined culture densities. Similar to the acute glucose shift experiment, *trr1Δ* cells exhibited a high frequency of focus formation during exponential phase that further increased in the aging culture (Fig. 6B and C). In contrast, no foci were observed in wild type cells until a threshold at approximately OD_600_=5, after which nearly all cells contained one or two large JUNQ foci. At three days of continuous culture, Hsp42-GFP localization in the two strains was virtually indistinguishable, suggesting that these phenomena are tightly linked and indicative of a common origin that connects redox maintenance with cellular energy levels.

**Figure 6.**
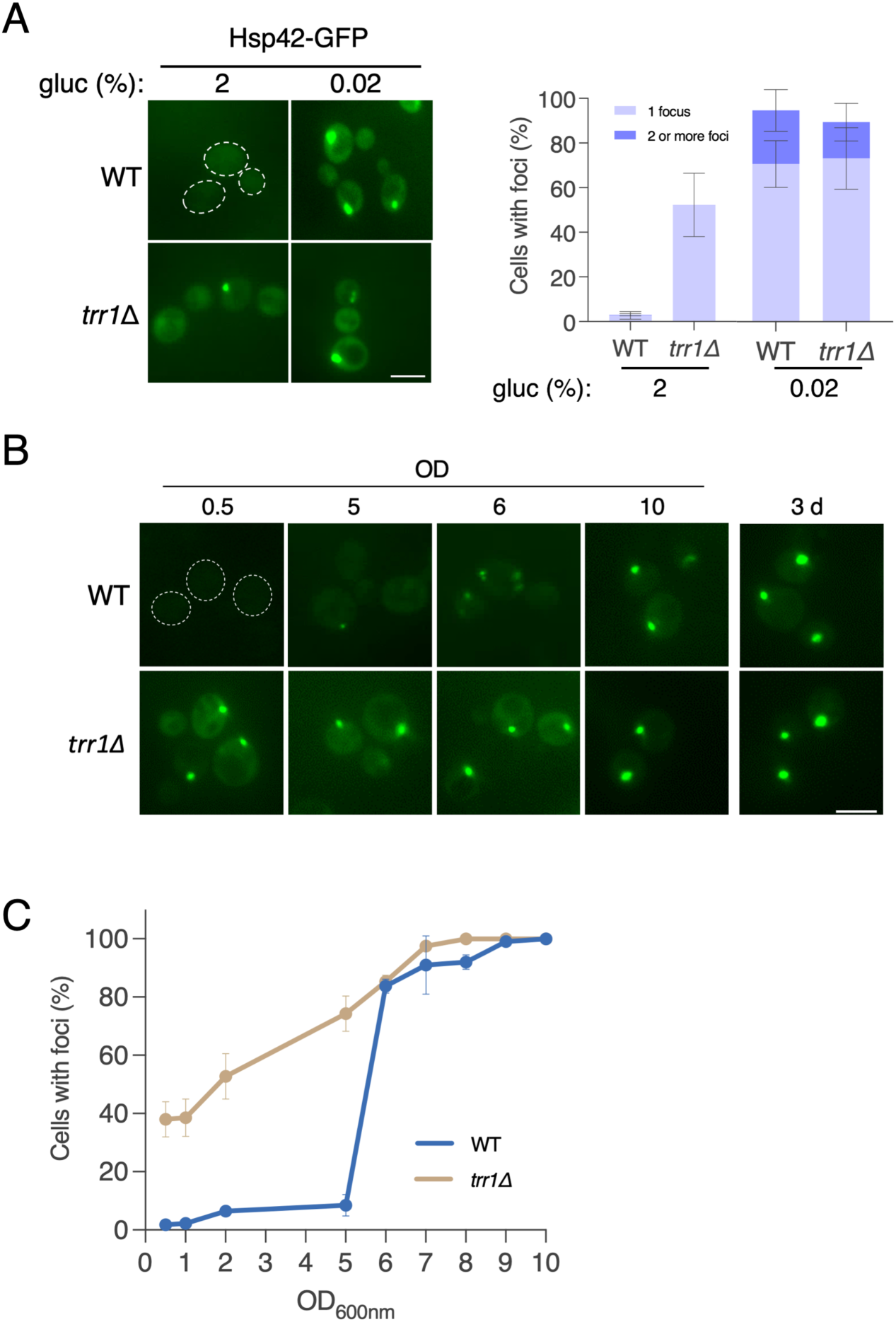
Hsp42 localization in glucose-starved and chronologically aging cells is phenocopied in *trr1Δ* cells. A) The indicated strains were grown at 30°C to mid-log phase and then either maintained in standard YPD medium or shifted to low glucose medium (YP + 0.02% glucose) for 90 min. The percentage of cells with one or two or more foci was determined by counting at least 100 cells from multiple fields. B) The same strains as in (A) were grown at 30°C and culture aliquots removed at the indicated optical densities (OD) or after 3 days of continuous growth. C) The percentage of cells with foci was determined by counting at least 100 cells from multiple fields. Scale bar = 5 µm.

## Discussion

Our work in redox-imbalanced cells has revealed a profound dysregulation of not only the protein misfolding-induced HSR, but also spatial quality control, manifest as specific and persistent hyperaccumulation of the cytosolic sequestrase Hsp42. How does loss of either thioredoxin reductase or the redundant cytosolic thioredoxins Trx1/2 lead to Hsf1 activation? Recent work has conclusively demonstrated that the Hsp70 chaperone proteins Ssa1/2 restrain Hsf1 in an inactive state through direct physical interaction at sites within the disordered amino- or carboxyl-terminal domains of the transcription factor (Zheng *et al*., 2016; Peffer *et al*., 2019; Chowdhary *et al*., 2021). The Ssa1-Hsf1 complex exhibits decreased DNA binding and nuclear condensate clustering, resulting in diminished RNA polymerase II recruitment at target gene promoters. Unfolded or misfolded polypeptides displaying exposed Hsp70 binding motifs compete with Hsf1 for Ssa1 binding, leading to HSR activation (Zheng *et al*., 2016). Protein misfolding due to redox imbalance could arise from defects in translation or multimerization/assembly of proteins containing one or more redox-active cysteine or methionine residues. However, we demonstrated that the heat-labile enzyme luciferase folds and acquires enzymatic activity in *trr1Δ* cells, arguing against a general protein folding defect. Alternatively, abridged function of the ubiquitin-proteasome pathway could generate a pool of misfolded proteins that compete for Hsf1-bound Ssa1 by blocking degradation, but our observation that bulk ubiquitination is unaffected in *trr1Δ* cells is inconsistent with this idea. We favor a model wherein impaired mobilization from CytoQ/JUNQ compartments results in a pool of accumulated misfolded proteins. This last scenario comports both with the persistent Hsp42 foci observed in *trr1Δ* cells and our findings that the unfoldable substrates CPY^‡^-GFP and tGnd1-GFP colocalized with these structures. Finally, it is possible that two independent mechanisms may be in play, as we have shown that reactive cysteines 264 and 303 in Ssa1 are subject to oxidative modification that inhibits general chaperone activity and releases Hsf1 (Santiago and Morano, 2022). Indeed, a pool of partially oxidized Ssa1 may exist in *trr1Δ* cells that both contributes to improper spatial quality control and improperly restrains Hsf1.

Notably, HSR activation was observed exclusively in mutant strains deficient in the cytosolic thioredoxin system. Elimination of other redox pathways, including the mitochondrial thioredoxin system, the cytosolic/vacuolar glutathione system, or the Mxr1 methionine-S-oxide reductase, did not result in elevated Hsf1 transcriptional activity. While the glutathione pathway plays a major role in redox balance in most eukaryotic cells, including humans, its importance in *Saccharomyces cerevisiae* is focused on detoxification, antioxidant function, amino acid biosynthesis and iron-sulfur cluster biogenesis, and may serve a subordinate role to the thioredoxin system. Interestingly, it was previously demonstrated that the endoplasmic reticulum-localized unfolded protein response pathway (UPR) is constitutively activated in *trr1Δ* cells, likely due to the shift toward oxidized thioredoxins in the absence of thioredoxin reductase, disallowing proper disulfide bond rearrangement by the protein disulfide isomerase Pdi1 (Kritsiligkou *et al*., 2018). In a similar manner, cytosolic protein cysteines could become oxidized in an environment lacking transferable reducing equivalents normally supplied by thioredoxins and the peroxiredoxin family. In support of this model, the peroxiredoxin Tsa1 localizes with oxidatively damaged, misfolded proteins (Hanzén *et al*., 2016). Importantly, we are not the first to observe dysregulation of the HSR in *trr1Δ* cells. MacDiarmid and coworkers identified a *TRR1*-inactivating genetic suppressor of *tsa1Δ* growth defects in cells deficient in zinc, and further showed that *trr1Δ* cells exhibited constitutive activation of the HSR that was exacerbated by zinc limitation (Macdiarmid *et al*., 2013). We have significantly extended these findings by identifying additional, previously unknown defects in spatial quality control in the absence of thioredoxin reductase in cells with no zinc challenge.

What is the nature of the exaggerated Hsp42-containing JUNQ compartment in *trr1Δ* cells? We found that other chaperones typically associated with CytoQ bodies and the JUNQ, such as Hsp40 (Ydj1), Hsp70 (Ssa1, Ssa4, Ssb1) and the disaggregase Hsp104, did not accumulate along with Hsp42 and misfolded protein. These chaperones transiently localize to CytoQ/JUNQ and participate in the extraction of polypeptides for further processing – refolding or proteasomal degradation. In fact, we found that Hsp104 and Hsp42 appropriately localized to CytoQ structures during acute heat shock. Hsp104 then appeared to return to a diffuse state while Hsp42 did not. Because large Hsp42 containing foci both preceded the heat shock experiment and persisted during recovery, we cannot distinguish whether the transient CytoQ-localized Hsp42 resolved like Hsp104 or became trapped within the accumulating JUNQ compartment. However, the impact of redox imbalance on spatial quality control seems specific to Hsp42 and likely associated misfolded proteins, rather than a general defect in protein processing. This point is underscored by our observation that the related sequestrase Btn2, associated with the INQ compartment, did not accumulate in detectable foci in *trr1Δ* cells, nor did the holdase Hsp26. Further work will be required to identify a mechanistic explanation for aberrant Hsp42 cellular dynamics. However, the importance of Hsp42 in cells experiencing redox imbalance through loss of Trr1 is underscored by the dramatic synergistic growth phenotype observed in *trr1Δhsp42Δ* cells. This growth defect is even more striking when considered in light of the fact that cells lacking *HSP42* exhibit little to no demonstrable phenotypes of any kind unless the Hsp70 system is compromised (Haslbeck *et al*., 2004; Ho *et al*., 2019). Clearly, Hsp42 is required to support proliferation of *trr1Δ* cells, which already experience profound growth retardation linked to ROS production in the ER. Our observation that *trr1Δhsp42Δ* growth rates are even further compromised during oxidative stress is consistent with a model wherein cytosolic sequestrase activity is critical in a challenging redox environment.

A notable feature of this work is the connection between our observations and previous studies identifying Hsp42 bodies forming in response to nutrient starvation and chronological aging. Indeed, it is difficult to distinguish Hsp42-GFP localization patterns under the various conditions. Our chronological aging experiment convincingly demonstrates that the exaggerated single foci that chronically persist in *trr1Δ* cells very closely resemble the same structures that begin to accumulate in post-logarithmic phase growth, which in turn resemble cells experiencing acute glucose starvation (0.02% glucose). While it is tempting to speculate that the observed nutrient starvation effects are therefore due to redox imbalance, we also note that the percentage of the *trr1Δ* cell population with Hsp42 foci increased during the aging experiment as well as upon shift to limiting glucose, suggestive of an additive relationship. Hsp42 localization patterns were not appreciably different in wild type or *trr1Δ* cells at 0.02% glucose, further suggesting that carbon source starvation (and hence energy production) is a more severe proteotoxic insult with respect to Hsp42/JUNQ dynamics than redox imbalance. Many questions remain to be answered regarding the links between redox homeostasis, energy state, and spatial quality control. Why is Hsp42 localization (JUNQ) so drastically altered with no detectable change in Btn2 dynamics (INQ) given the interconnectedness of the two compartments (Sontag *et al*., 2023)? What proteins constitute the exaggerated JUNQ compartment present in *trr1Δ* and aging/starved cells, and are they similar among the different conditions? Finally, we seek to understand the cytoprotective role that Hsp42 appears to play in redox-challenged cells, presumably via sequestration of potentially toxic misfolded proteins.

## Materials and Methods

### Strains and plasmids

All strains used in this study are isogenic to BY4741 and are listed in Table 1. Multiple GFP fusion strains were from the ThermoFisher GFP Collection (Huh *et al*., 2003). Multiple gene deletion strains are from the Yeast Knockout Collection (Winzeler *et al*., 1999). Construction of knockout strains was done by generating PCR amplicons containing one of the markers G418^R^, *LEU2* or *URA3,* flanked by upstream and downstream noncoding regions of the knockout target gene. Alternatively, we employed the seamless deletion method to recycle the *URA3* gene and generate *trr1*Δ seamless strains, denoted as *trr1Δ\** in Table 1 (Horecka and Davis, 2014). For expression of *TRR1* in *trr1*Δ strains, the genomic region of BY4741 containing the *TRR1* coding sequence and 1 kb upstream and 1 kb downstream was amplified and cloned into the pRS415 plasmid (Table 1) using introduced *SpeI* and *XbaI* restriction sites. The same plasmid construct was used for the generation of *trr1-C142S C145S*, where the codons encoding the two cysteines in the catalytic region of Trr1 (C142 and C145) were changed to encode serine using overlap PCR. The GFP-expressing control plasmid pRS415-GPD-GFP was constructed by amplifying the GFP coding sequence from the genomic DNA of the Hsp42-GFP strain and cloned into pRS415-GPD using introduced *SpeI* and *XhoI* restriction sites. All plasmid constructions and genomic integration cassettes were verified by sequencing prior to yeast transformation. Yeast transformation was performed using the rapid yeast transformation protocol (Gietz *et al*., 1992). Plasmids pRH2081 (TDH3-CPY^‡^-GFP), pRH2476 (TDH3-tGND-GFP) and pRS303-*PHO88-mCherry* were linearized as previously described before integrative transformation (Heck *et al*., 2010; D’Urso *et al*., 2016).

**Table 1.**
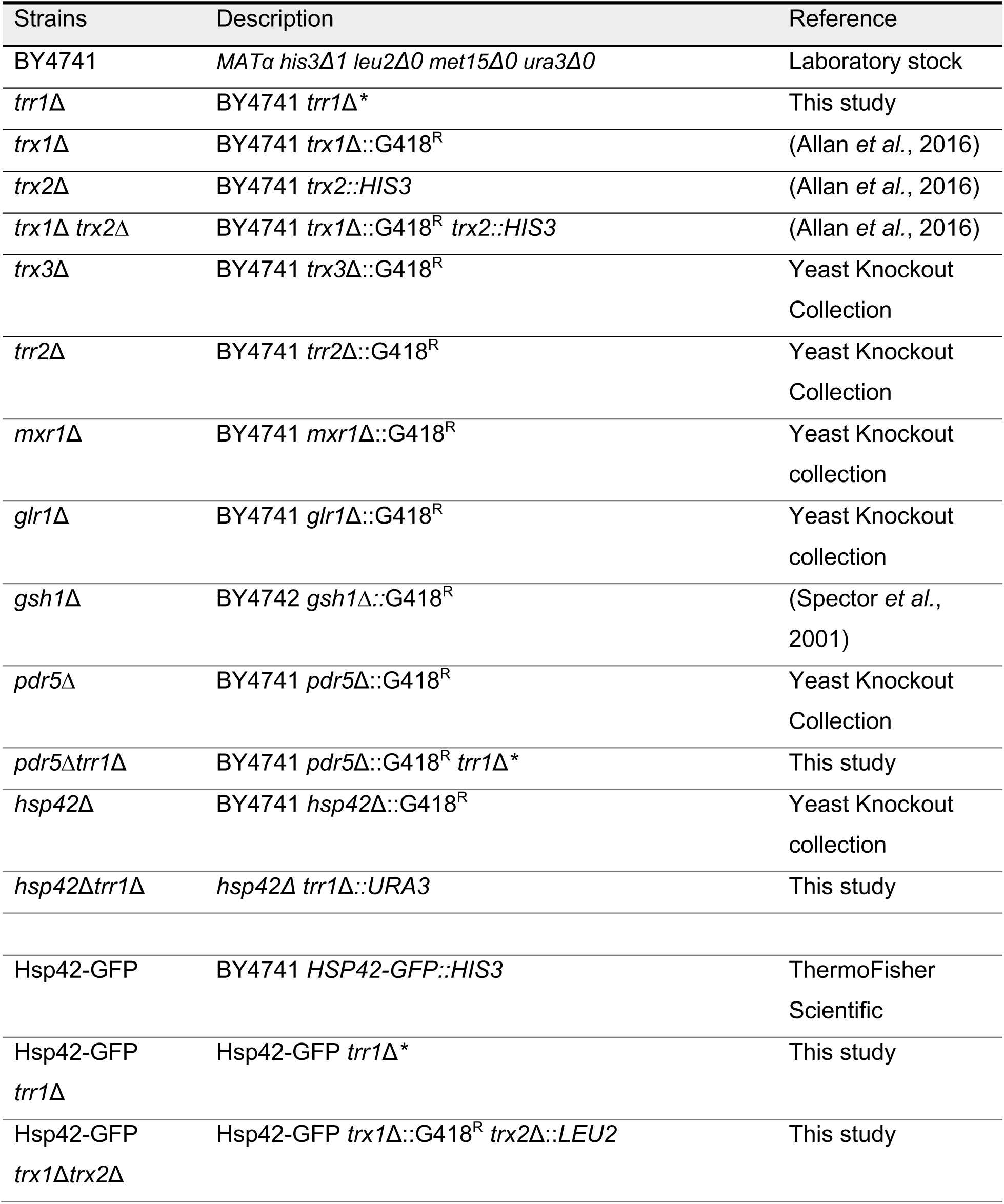
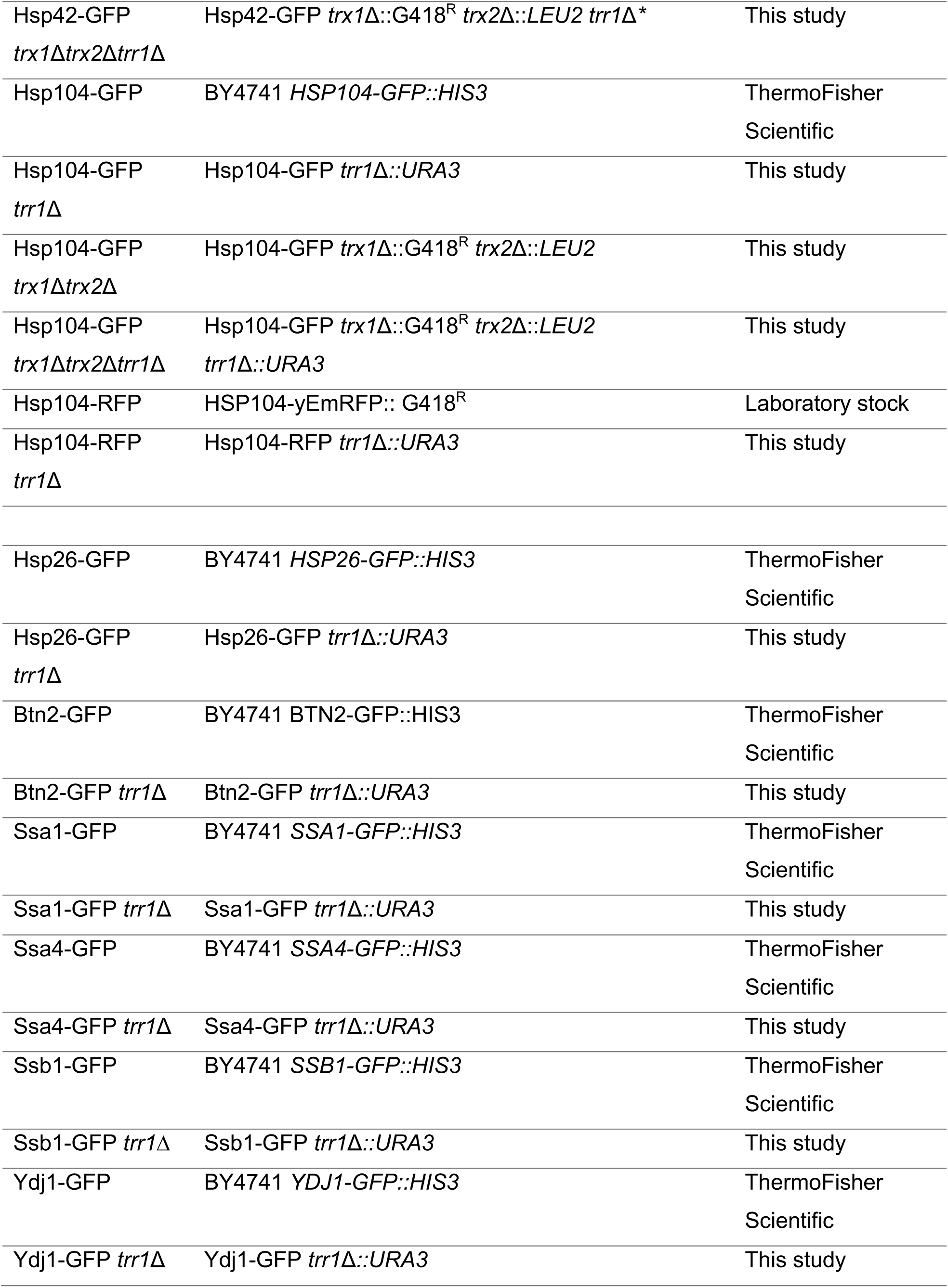
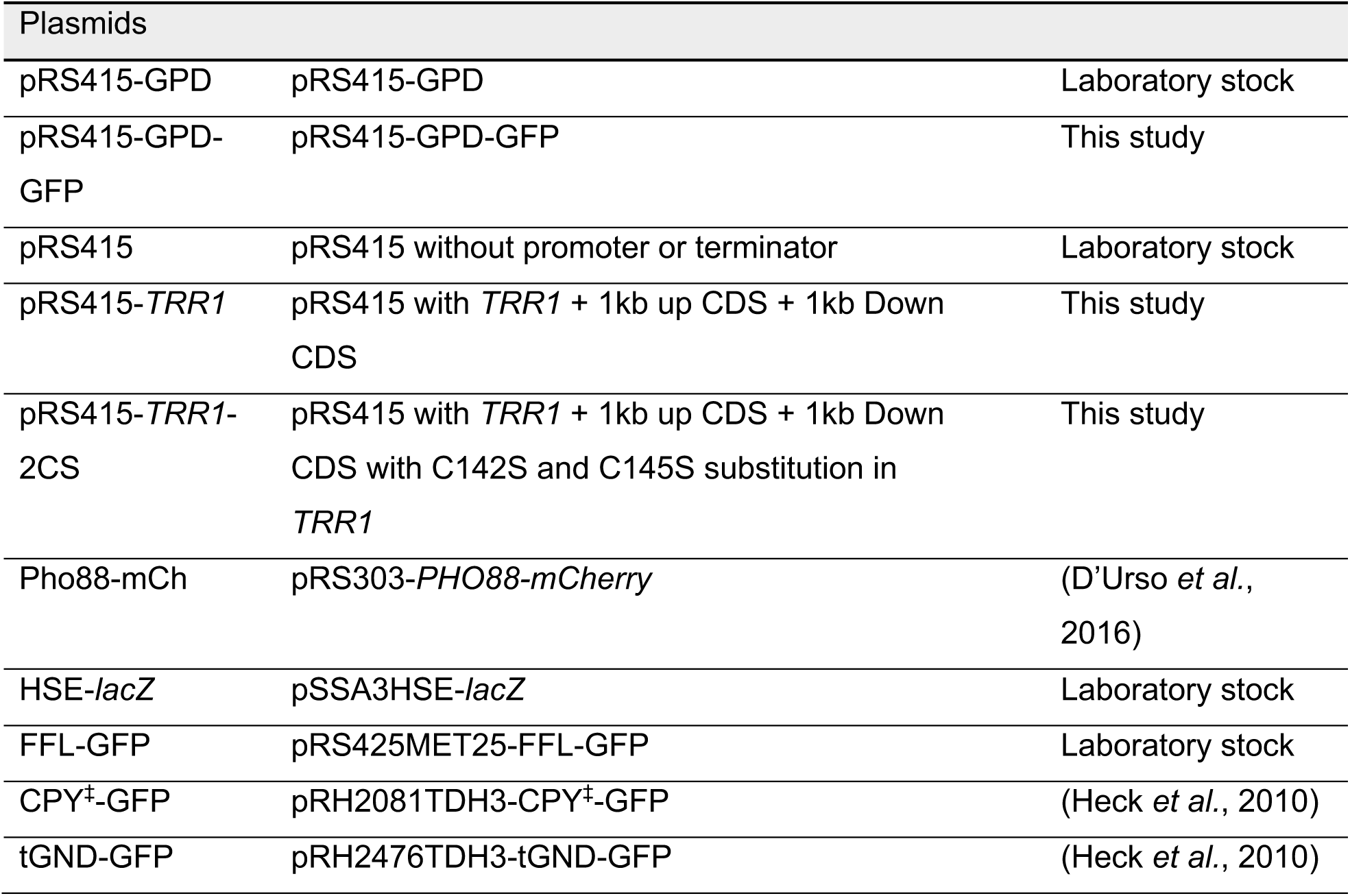
Yeast strains and plasmids

### Yeast culture and growth assays

Strains were cultured in standard non-selective (YPD; 1% yeast extract, 2% peptone, 2% glucose) medium or selective synthetic complete medium (SC; 2% glucose) lacking amino acids for marker selection (Sunrise Science). Strains were grown at 30° C with aeration to mid-log phase unless otherwise specified. For growth curve assays, mid-log phase cells were inoculated at a low culture density (OD_600_ = 0.01) in 100 µl fresh medium in a 96-well sterile plate and incubated with intermittent agitation in a Synergy MX Microplate reader (Biotek Instruments) at 30° C. Heat shock experiments were performed in a shaking waterbath at 39°C in glass tubes or flasks for 20 min, unless otherwise specified. For the cycloheximide (CHX) chase experiments, CHX was added at 100 µg/ml final concentration for indicated times. To monitor Hsp42-GFP dynamics during glucose starvation, mid-log phase cells were transferred to 0.02% glucose YP medium for 90 min. Proteasome inhibition was achieved via addition of 75 µM carbobenzoxy-Leu-Leu-leucinal (MG-132, MilliporeSigma) in strains additionally carrying the *pdr5Δ* deletion to allow for uptake (Liu et al., 2007)(Collins *et al*., 2010).

### Fluorescence microscopy

Cells were wet-mounted on glass slides and imaged immediately using an Olympus IX81-ZDC inverted microscope with a 100x objective lens using fluorescence with appropriate standard filter sets. Images were captured with a Hamamatsu ORCA camera. Quantitation was done by counting at least 100 cells and dividing the number of cells containing aggregates by the total number of cells counted. For some experiments, cells containing a single, large focus were distinguished from those containing two or more visible foci.

### HSR activation using HSE*-lacZ* reporter

Activity of Hsf1 was determined by adding 50 µl of cell suspension (cells expressing the pSSA3HSE-*lacZ* plasmid) to 50 µl of β-Glo reagent (Promega) in a white 96-well plate. After 30 min of incubation at 30°C, luminescence was measured in a Synergy MX Microplate reader (Abrams et al., 2014).

### In vivo firefly luciferase refolding assay

Refolding of the heat-labile enzyme firefly luciferase was performed as described with the following modifications (Abrams and Morano, 2013). Cells bearing the plasmid p425MET25-FFL-GFP-leu2::URA3 were grown to mid-log phase in selective medium supplemented with 1 mM methionine at 30°C. Cells were washed and sub-cultured in medium lacking methionine to induce expression of FFL-GFP from the *MET25* promoter for 1 hr. Cycloheximide was added to cultures at 100 µg/ml final concentration to halt protein synthesis. Basal FFL activity was determined by adding 10 µl of 222 nM luciferin to 100 µl of cell culture and measuring light production using a Synergy MX Microplate reader equipped with a luminometer. Cultures were shifted to 42°C for 15 min to allow heat denaturation of the FFL enzyme, and FFL activity measured after return to 30°C growth conditions for multiple time points during the recovery period.

### Proteasomal activity

Proteasomal activity was assessed using the Proteasome 20S Activity Assay Kit (MAK172-1KT; MilliporeSigma). Experiments were performed by adding 50 µl of cell suspension to 50 µl of required reagent in a white 96-well plate. After 2 hr incubation at 30° C, 20S proteasome activity was measured in a Synergy MX Microplate reader.

### Western blots

Proteins were isolated using glass bead lysis as described (Abrams et al, 2014). Protein samples were heated (65°C, 15 min) in SDS-PAGE sample-loading buffer, loaded into 10% bis-acrylamide/SDS gels and separated by electrophoresis. Separated protein samples were transferred to a polyvinylidene difluoride (PVDF) membrane, blocked in 5% nonfat dry milk and proteins detected using monoclonal anti-GFP (Roche), monoclonal anti-PGK (Invitrogen), and monoclonal anti-Ub (EMD Millipore) antibodies, followed by detection with HRP-conjugated secondary antibody (MilliporeSigma) with enhanced chemiluminescence reagent and direct photon emission imaging. Protein bands were quantitated using Image Studio Lite (LI-COR Biosciences).

### RNA isolation and qRT-PCR

Cultures were grown in 20 ml of YPD medium at 30°C, harvested at OD_600_ 0.8, centrifuged, and the cell pellet immediately frozen on dry ice. Total RNA was isolated by the hot phenol method. For qRT-PCR assays, 1 µg of RNA was converted to cDNA using the iScript cDNA synthesis kit (Bio-Rad). Relative expression of the *SSA3* and *SSA4* genes was measured by qRT-PCR using iTaq Universal SYBR Green Supermix (Bio-Rad) and calculated using standard method (Nolan *et al*., 2006). *TAF10* was used as the normalization control gene. All experiments were conducted with three technical and three biological replicates.

### Statistical analysis

Significance was determined using GraphPad QuickCalcs Welch’s unpaired *t* test calculator (GraphPad, Dotmetrics v. 9.3.0).

## Abbreviations

sHSP: small heat shock protein
HSR: heat shock response
INQ: internuclear quality control compartment
JUNQ: juxtanuclear quality control compartment
IPOD: insoluble protein deposit
UPR: unfolded protein response
UPS: ubiquitin-proteasome system

## Acknowledgements

This work was supported by grants R01GM127287 and R35GM149196 from the National Institutes of Health. We thank Drs. Jason Brickner (Northwestern University), Randy Hampton (UC San Diego), Michel Toledano (Université Paris-Saclay) and James West (College of Wooster) for reagents and James West for helpful discussions and critical reading of the manuscript.

## Supplemental Information

**Figure S1.**
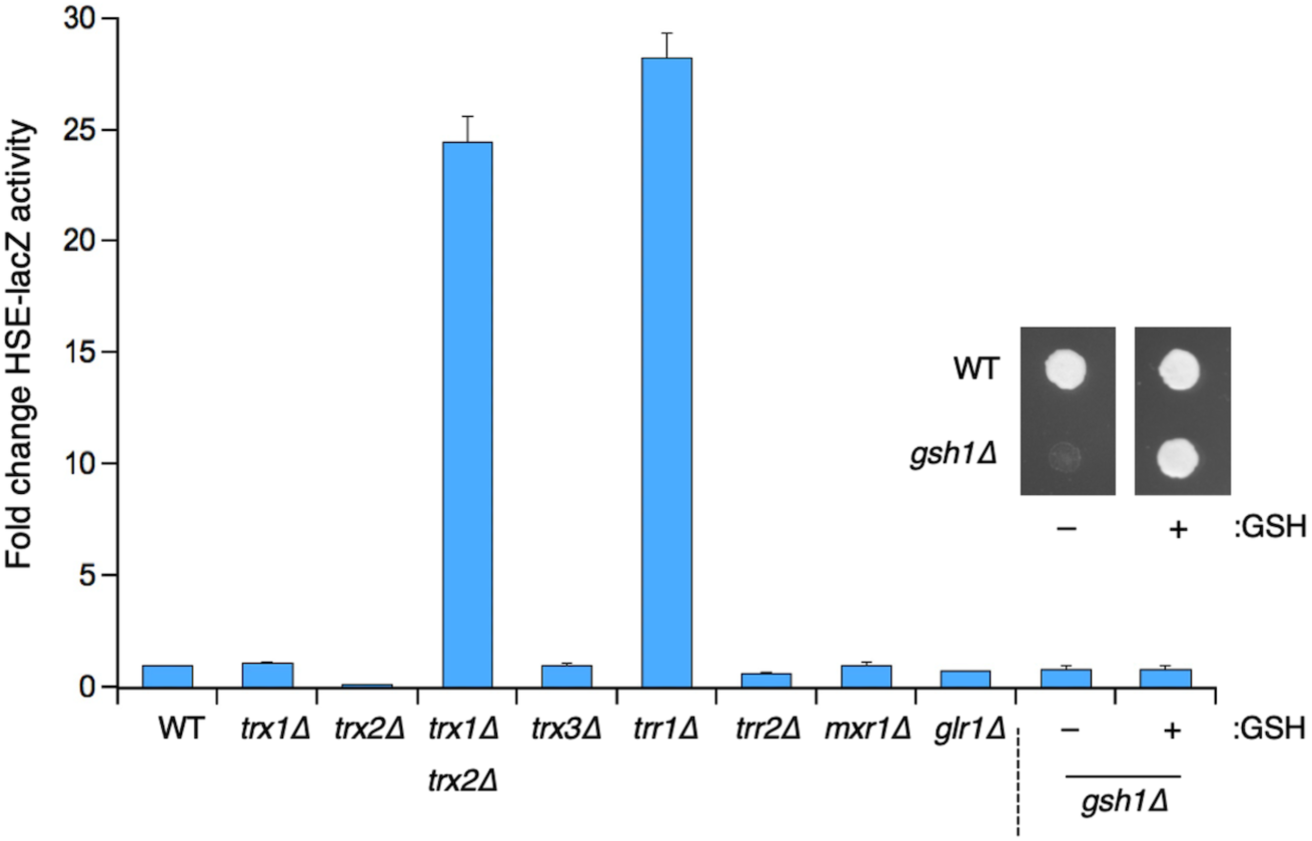
Chronic activation of the heat shock response occurs exclusively in cells lacking the cytosolic thioredoxin system. The indicated strains with null mutations in genes comprising different redox pathways were transformed with the pSSA3HSE-lacZ plasmid and basal β-galactosidase activity determined as described in Materials and Methods. Values are mean fold activation relative to wild type with error bars denoting standard deviation. To assess the HSR in *gsh1Δ* cells, reduced glutathione was added to cultures grown to mod-log phase, after which the cells were washed and diluted into glutathione-free medium and grown for an additional 6 hr prior to β-galactosidase activity determination. Inset: 3 µl aliquots of the indicated cultures were spotted onto solid medium lacking or containing 1 mM glutathione.

**Figure S2:**
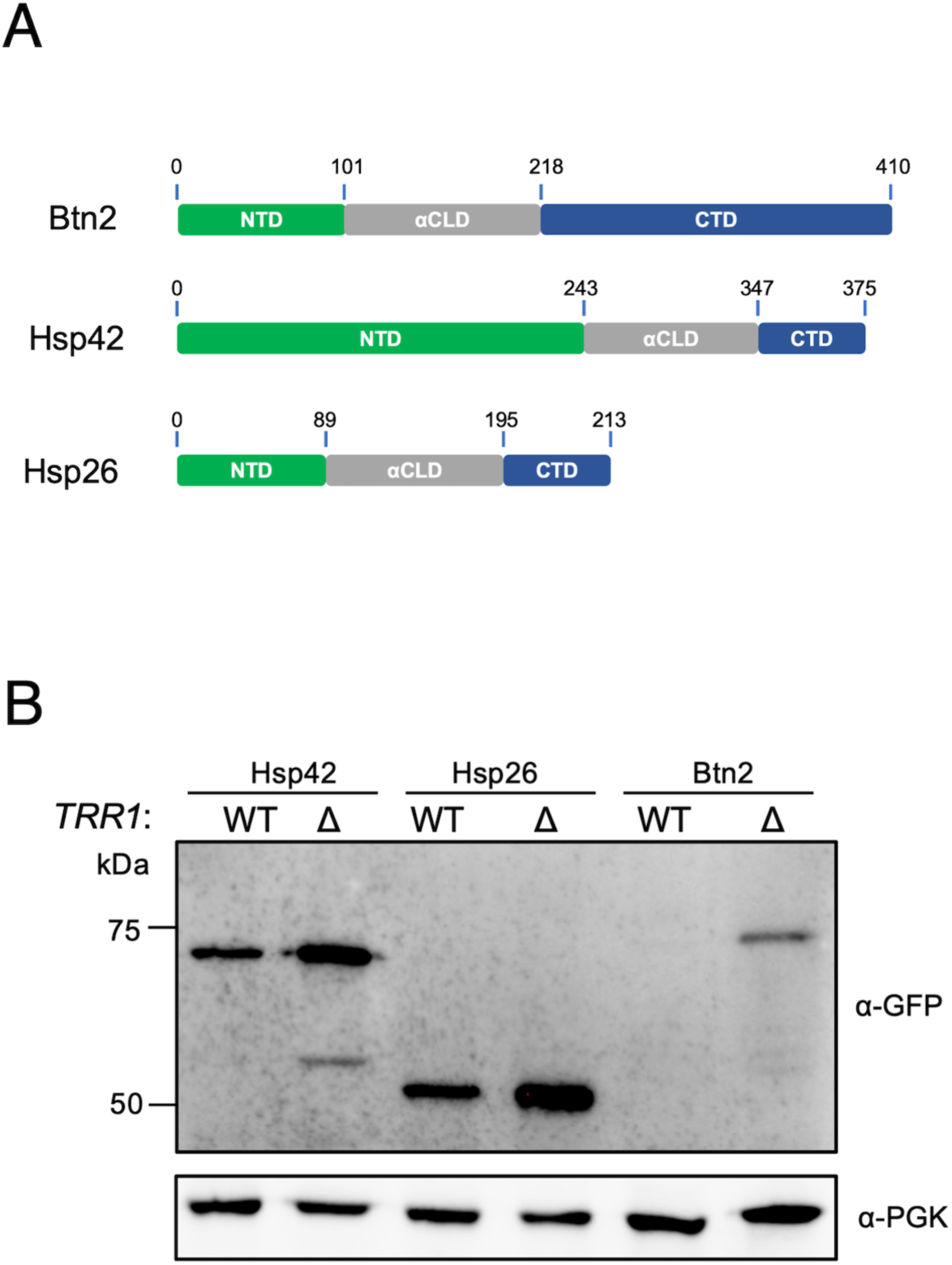
Expression of all three known yeast small heat shock proteins is elevated in *trr1Δ* cells. A) Domain schematics of the indicated proteins. NTD, amino terminal domain; αCLD, α-crystallin-like domain; CTD, carboxyl terminal domain. B) SDS-PAGE immunoblot of protein extracts prepared from the indicated strains bearing chromosomal GFP fusions grown to mid-log phase.

**Figure S3:**
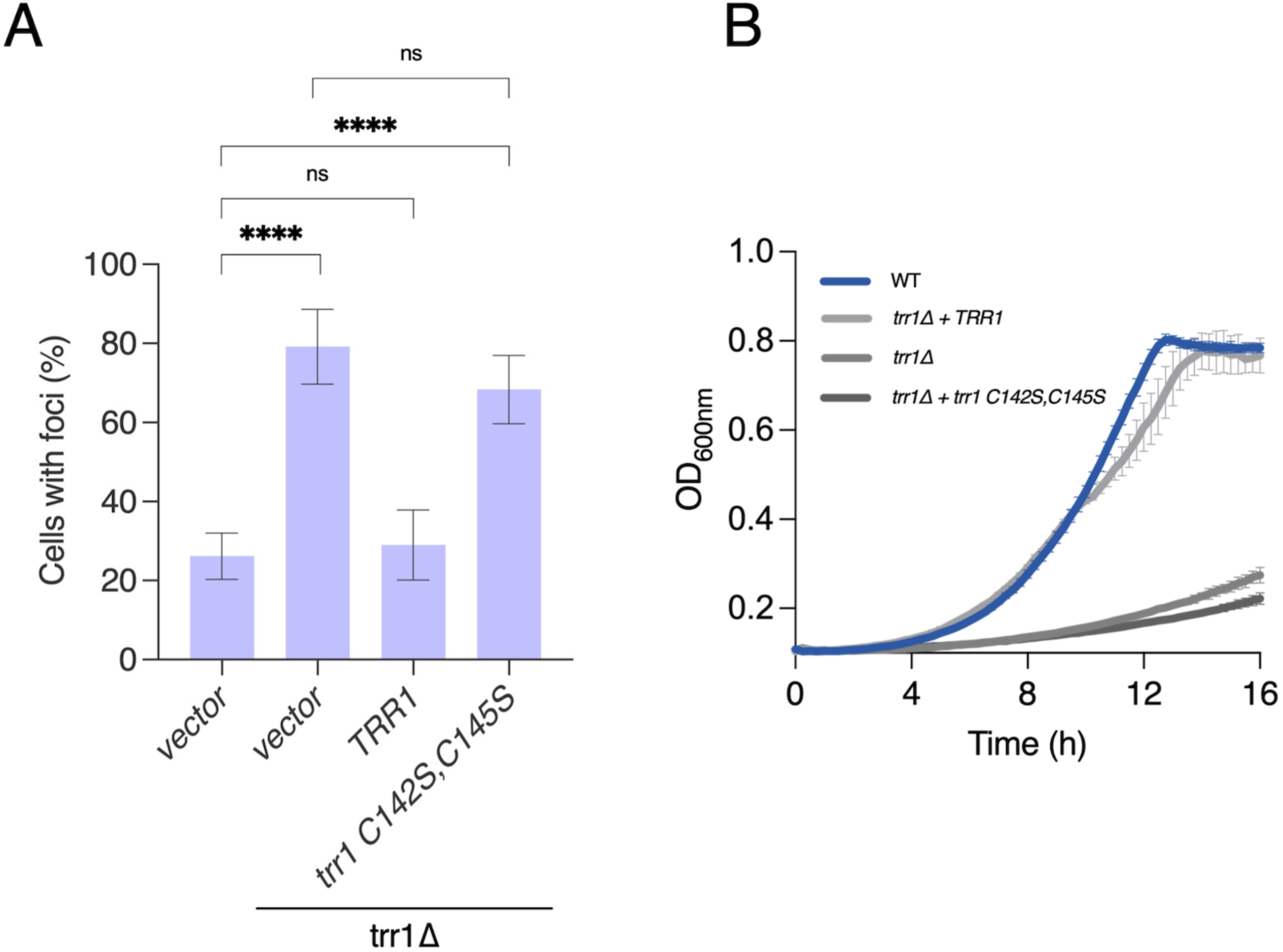
Thioredoxin reductase activity of Trr1 is required for normal Hsp42 spatial distribution. A) The indicated cells (WT with vector alone, or *trr1Δ* cells bearing the indicated plasmids) were grown to mid-log phase and the percentage of cells with foci was determined by counting at least 100 cells from multiple fields. B) The same strains as in (A) were inoculated at an initial OD_600_=0.01 in a sterile 96-well plate and grown with shaking at 30°C for 16 h with density measurements taken every 10 min. The average of three biological replicate growth curves are shown with standard deviation. Statistical significance between the indicated strains was determined using Welch’s unpaired t test (p=0.05, *; p=0.005, **; p=0.0005, ***; p=0.00005, ****).

**Figure S4:**
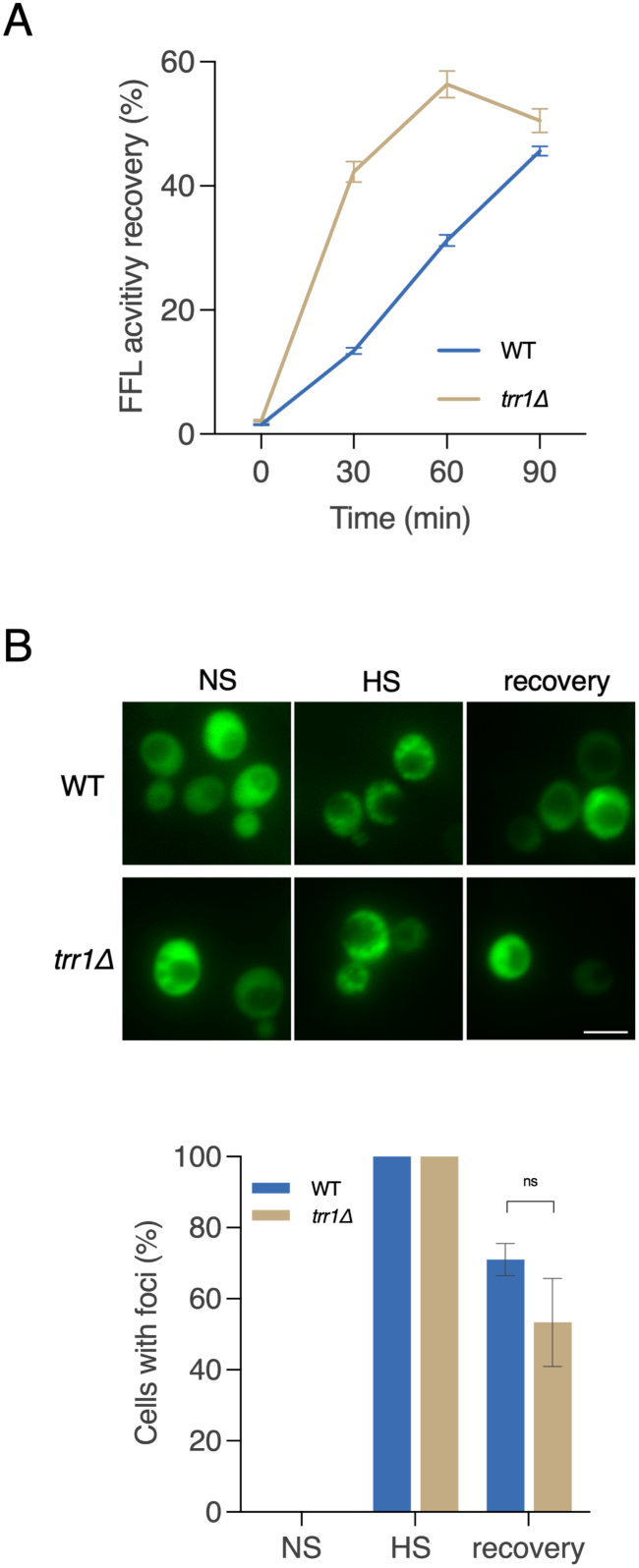
Folding of the model protein luciferase is not impaired in *trr1Δ* cells. A) WT or *trr1Δ* cells transformed with the FFL-GFP plasmid were grown to mid-log phase and steady state FFL activity measured as described in Materials and Methods. Cells were then heat shocked at 42°C for 15 min to cause FFL unfolding. FFL activity in living cells was measured at the indicated time points after return to growth at 30°C. B) Aliquots of cells from (A) were removed and imaged to identify FFL-GFP CytoQ aggregates. C) The percentage of cells in (B) with foci was determined by counting at least 100 cells from multiple fields. Scale bar = 5 µm. Statistical significance between the indicated strains was determined using Welch’s unpaired t test (p=0.05, *; p=0.005, **; p=0.0005, ***; p=0.00005, ****).

**Figure S5:**
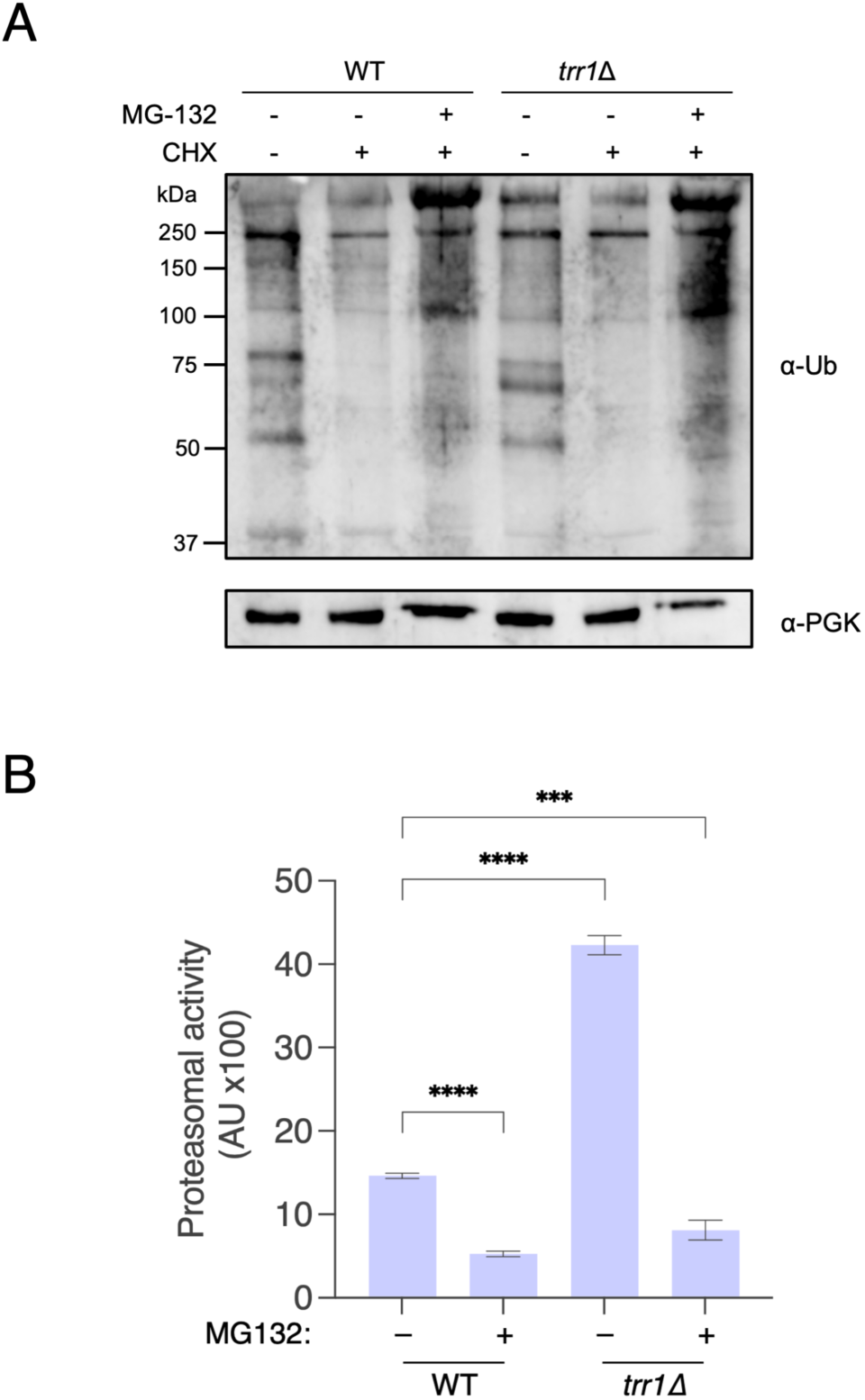
The ubiquitin-proteasome system is not impaired in *trr1Δ* cells. A) WT and *trr1Δ* cells (also bearing the *pdr5Δ* mutation) were grown to mid-log phase and treated or not with MG-132 at 75 µM or cycloheximide (CHX, 100 µg/ml) for 2 hr. Protein extracts were prepared, resolved by SDS-PAGE and ubiquitin profiles determined by blotting using anti-ubiquitin antibody. Levels of PGK1 were determined as a load control. B) The same strains as in (A) were grown to mid-log phase and MG-132-sensitive 20S proteasomal activity determined as described in Materials and Methods. Statistical significance between the indicated strains was determined using Welch’s unpaired t test (p=0.05, *; p=0.005, **; p=0.0005, ***; p=0.00005, ****).

## References

1. Abrams, JL, and Morano, KA (2013). Coupled assays for monitoring protein refolding in Saccharomyces cerevisiae. J Vis Exp.

2. Akerfelt, M, Morimoto, RI, and Sistonen, L (2010). Heat shock factors: integrators of cell stress, development and lifespan. Nat Rev Mol Cell Biol 11, 545–555.

3. Allan, KM et al. (2016). Trapping redox partnerships in oxidant-sensitive proteins with a small, thiol-reactive cross-linker. Free Radic Biol Med 101, 356–366.

4. Ayer, A, Gourlay, CW, and Dawes, IW (2014). Cellular redox homeostasis, reactive oxygen species and replicative ageing in Saccharomyces cerevisiae. FEMS Yeast Res 14, 60–72.

5. Baker, HA, and Bernardini, JP (2021). It’s not just a phase; ubiquitination in cytosolic protein quality control. Biochem Soc Trans 49, 365–377.

6. Bonner, JJ, Ballou, C, and Fackenthal, DL (1994). Interactions between DNA-bound trimers of the yeast heat shock factor. Mol Cell Biol 14, 501–508.

7. Chae, HZ, Chung, SJ, and Rhee, SG (1994). Thioredoxin-dependent peroxide reductase from yeast. J Biol Chem 269, 27670–27678.

8. Chowdhary, S, Kainth, AS, Paracha, S, Gross, DS, and Pincus, D (2021). Inducible transcriptional condensates drive 3D genome reorganization in the heat shock response. BioRxiv, 2021.10.27.466170.

9. Collins, GA, Gomez, TA, Deshaies, RJ, and Tansey, WP (2010). Combined chemical and genetic approach to inhibit proteolysis by the proteasome. Yeast Chichester Engl 27, 965–974.

10. Dahl, JU, Gray, MJ, and Jakob, U (2015). Protein quality control under oxidative stress conditions. J Mol Biol 427, 1549–1563.

11. D’Urso, A, Takahashi, Y-H, Xiong, B, Marone, J, Coukos, R, Randise-Hinchliff, C, Wang, J-P, Shilatifard, A, and Brickner, JH (2016). Set1/COMPASS and Mediator are repurposed to promote epigenetic transcriptional memory. ELife 5, e16691.

12. Escusa-Toret, S, Vonk, WIM, and Frydman, J (2013). Spatial sequestration of misfolded proteins by a dynamic chaperone pathway enhances cellular fitness during stress. Nat Cell Biol 15, 1231–1243.

13. Fra, A, Yoboue, ED, and Sitia, R (2017). Cysteines as Redox Molecular Switches and Targets of Disease. Front Mol Neurosci 10.

14. Gietz, D, St Jean, A, Woods, RA, and Schiestl, RH (1992). Improved method for high efficiency transformation of intact yeast cells. Nucleic Acids Res 20, 1425.

15. Grousl, T, Ungelenk, S, Miller, S, Ho, CT, Khokhrina, M, Mayer, MP, Bukau, B, and Mogk, A (2018). A prion-like domain in Hsp42 drives chaperonefacilitated aggregation of misfolded proteins. J Cell Biol 217, 1269–1285.

16. Hahn, J-S, Neef, DW, and Thiele, DJ (2006). A stress regulatory network for co-ordinated activation of proteasome expression mediated by yeast heat shock transcription factor. Mol Microbiol 60, 240–251.

17. Hanzén, S et al. (2016). Lifespan Control by Redox-Dependent Recruitment of Chaperones to Misfolded Proteins. Cell 166, 140–151.

18. Haslbeck, M, Braun, N, Stromer, T, Richter, B, Model, N, Weinkauf, S, and Buchner, J (2004). Hsp42 is the general small heat shock protein in the cytosol of Saccharomyces cerevisiae. EMBO J 23, 638–649.

19. Heck, JW, Cheung, SK, and Hampton, RY (2010). Cytoplasmic protein quality control degradation mediated by parallel actions of the E3 ubiquitin ligases Ubr1 and San1. Proc Natl Acad Sci U S A 107, 1106–1111.

20. Herman, PK (2002). Stationary phase in yeast. Curr Opin Microbiol 5, 602–607.

21. Hill, SM, Hanzén, S, and Nyström, T (2017). Restricted access: spatial sequestration of damaged proteins during stress and aging. EMBO Rep 18, 377–391.

22. Ho, C et al. (2019). Cellular sequestrases maintain basal Hsp70 capacity ensuring balanced proteostasis. Nat Commun 10, 1–15.

23. Horecka, J, and Davis, RW (2014). The 50:50 method for PCR-based seamless genome editing in yeast. Yeast Chichester Engl 31, 103–112.

24. Huh, WK, Falvo, JV, Gerke, LC, Carroll, AS, Howson, RW, Weissman, JS, and O’Shea, EK (2003). Global analysis of protein localization in budding yeast. Nature 425, 686–691.

25. Kaganovich, D, Kopito, R, and Frydman, J (2008). Misfolded proteins partition between two distinct quality control compartments. Nature 454, 1088–1095.

26. Kaya, A, Koc, A, Lee, BC, Fomenko, DE, Rederstorff, M, Krol, A, Lescure, A, and Gladyshev, VN (2010). Compartmentalization and regulation of mitochondrial function by methionine sulfoxide reductases in yeast. Biochemistry 49, 8618–8625.

27. Kritsiligkou, P, Rand, JD, Weids, AJ, Wang, X, Kershaw, CJ, and Grant, CM (2018). Endoplasmic reticulum (ER) stress–induced reactive oxygen species (ROS) are detrimental for the fitness of a thioredoxin reductase mutant. J Biol Chem 293, 11984– 11995.

28. Kumar, A, Mathew, V, and Stirling, PC (2022). Nuclear protein quality control in yeast: The latest INQuiries. J Biol Chem 298, 102199.

29. Le Moan, N, Clement, G, Le Maout, S, Tacnet, F, and Toledano, MB (2006). The Saccharomyces cerevisiae proteome of oxidized protein thiols: Contrasted functions for the thioredoxin and glutathione pathways. J Biol Chem 281, 10420–10430.

30. Lee, HY, Cheng, KY, Chao, JC, and Leu, JY (2016). Differentiated cytoplasmic granule formation in quiescent and non-quiescent cells upon chronological aging. Microb Cell 3, 109–119.

31. Lévy, E, Banna, NE, Baïlle, D, Heneman-Masurel, A, Truchet, S, Rezaei, H, Huang, ME, Béringue, V, Martin, D, and Vernis, L (2019). Causative links between protein aggregation and oxidative stress: A review. Int J Mol Sci 20.

32. Liu, IC, Chiu, SW, Lee, HY, and Leu, JY (2012). The histone deacetylase Hos2 forms an Hsp42-dependent cytoplasmic granule in quiescent yeast cells. Mol Biol Cell 23, 1231– 1242.

33. López-Mirabal, HR, and Winther, JR (2008). Redox characteristics of the eukaryotic cytosol. Biochim Biophys Acta - Mol Cell Res 1783, 629–640.

34. Macdiarmid, CW, Taggart, J, Kerdsomboon, K, Kubisiak, M, Panascharoen, S, Schelble, K, and Eide, DJ (2013). Peroxiredoxin chaperone activity is critical for protein homeostasis in zinc-deficient yeast. J Biol Chem 288, 31313–31327.

35. Malinovska, L, Kroschwald, S, Munder, MC, Richter, D, and Alberti, S (2012). Molecular chaperones and stress-inducible protein-sorting factors coordinate the spatiotemporal distribution of protein aggregates. Mol Biol Cell 23, 3041–3056.

36. Mathew, V, Tam, AS, Milbury, KL, Hofmann, AK, Hughes, CS, Morin, GB, Loewen, CJR, and Stirling, PC (2017). Selective aggregation of the splicing factor Hsh155 suppresses splicing upon genotoxic stress. J Cell Biol 216, 4027–4040.

37. Miller, SB et al. (2015a). Compartment-specific aggregases direct distinct nuclear and cytoplasmic aggregate deposition. EMBO J 34, 778–797.

38. Miller, SBM, Mogk, A, and Bukau, B (2015b). Spatially organized aggregation of misfolded proteins as cellular stress defense strategy. J Mol Biol 427, 1564–1574.

39. Miyata, Y, Rauch, JN, Jinwal, UK, Thompson, AD, Srinivasan, S, Dickey, CA, and Gestwicki, JE (2012). Cysteine reactivity distinguishes redox sensing by the heat inducible and constitutive forms of heat shock protein 70. Chem Biol 19, 1391–1399.

40. Mogk, A, and Bukau, B (2017). Role of sHsps in organizing cytosolic protein aggregation and disaggregation. Cell Stress Chaperones 22, 493–502.

41. Mogk, A, Ruger-Herreros, C, and Bukau, B (2019). Cellular functions and mechanisms of action of small heat shock proteins. Annu Rev Microbiol 73, 89–110.

42. Morano, KA, Santoro, N, Koch, KA, and Thiele, DJ (1999). A trans-Activation Domain in Yeast Heat Shock Transcription Factor Is Essential for Cell Cycle Progression during Stress. Mol Cell Biol 19, 402–411.

43. Narayanaswamy, R, Levy, M, Tsechansky, M, Stovall, GM, O’Connell, JD, Mirrielees, J, Ellington, AD, and Marcotte, EM (2009). Widespread reorganization of metabolic enzymes into reversible assemblies upon nutrient starvation. Proc Natl Acad Sci U S A 106, 10147–10152.

44. Nolan, T, Hands, RE, and Bustin, SA (2006). Quantification of mRNA using real-time RT-PCR. Nat Protoc 1, 1559–1582.

45. O’Connell, JD, Tsechansky, M, Royal, A, Boutz, DR, Ellington, AD, and Marcotte, EM (2014). A proteomic survey of widespread protein aggregation in yeast. Mol Biosyst 10, 851–861.

46. Peffer, S, Gonçalves, D, and Morano, KA (2019). Regulation of the Hsf1-dependent transcriptome via conserved bipartite contacts with Hsp70 promotes survival in yeast. J Biol Chem 294, 12191–12202.

47. Rothe, S, Prakash, A, and Tyedmers, J (2018). The Insoluble Protein Deposit (IPOD) in Yeast. Front Mol Neurosci 11.

48. Saarikangas, J, and Barral, Y (2015). Protein aggregates are associated with replicative aging without compromising protein quality control. ELife 4.

49. Santiago, A, and Morano, KA (2022). Oxidation of two cysteines within yeast Hsp70 impairs proteostasis while directly triggering an Hsf1-dependent cytoprotective response. J Biol Chem 298, 102424.

50. Sathyanarayanan, U, Musa, M, Bou Dib, P, Raimundo, N, Milosevic, I, and Krisko, A (2020). ATP hydrolysis by yeast Hsp104 determines protein aggregate dissolution and size in vivo. Nat Commun 11.

51. Shrivastava, A, Sandhof, CA, Reinle, K, Jawed, A, Ruger-Herreros, C, Schwarz, D, Creamer, D, Nussbaum-Krammer, C, Mogk, A, and Bukau, B (2022). The cytoprotective sequestration activity of small heat shock proteins is evolutionarily conserved. J Cell Biol 221.

52. Solís, EJ, Pandey, JP, Zheng, X, Jin, DX, Gupta, PB, Airoldi, EM, Pincus, D, and Denic, V (2016). Defining the Essential Function of Yeast Hsf1 Reveals a Compact Transcriptional Program for Maintaining Eukaryotic Proteostasis. Mol Cell 63, 60–71.

53. Sontag, EM, Morales-Polanco, F, Chen, J-H, McDermott, G, Dolan, PT, Gestaut, D, Le Gros, MA, Larabell, C, and Frydman, J (2023). Nuclear and cytoplasmic spatial protein quality control is coordinated by nuclear-vacuolar junctions and perinuclear ESCRT. Nat Cell Biol 25, 699–713.

54. Specht, S, Miller, SBM, Mogk, A, and Bukau, B (2011). Hsp42 is required for sequestration of protein aggregates into deposition sites in Saccharomyces cerevisiae. J Cell Biol 195, 617–629.

55. Spector, D, Labarre, J, and Toledano, MB (2001). A genetic investigation of the essential role of glutathione: mutations in the proline biosynthesis pathway are the only suppressors of glutathione auxotrophy in yeast. J Biol Chem 276, 7011–7016.

56. Tkach, JM, and Glover, JR (2008). Nucleocytoplasmic Trafficking of the Molecular Chaperone Hsp104 in Unstressed and Heat-Shocked Cells. Traffic 9, 39–56.

57. Trott, A, West, JD, Klaić, L, Westerheide, SD, Silverman, RB, Morimoto, RI, and Morano, KA (2008). Activation of heat shock and antioxidant responses by the natural product celastrol: Transcriptional signatures of a thiol-targeted molecule. Mol Biol Cell 19, 1104– 1112.

58. Trotter, EW, and Grant, CM (2003). Non-reciprocal regulation of the redox state of the glutathione-glutaredoxin and thioredoxin systems. EMBO Rep 4, 184–188.

59. Trotter, EW, and Grant, CM (2005). Overlapping roles of the cytoplasmic and mitochondrial redox regulatory systems in the yeast Saccharomyces cerevisiae. Eukaryot Cell 4, 392–400.

60. Trotter, EW, Kao, CM-F, Berenfeld, L, Botstein, D, Petsko, GA, and Gray, JV (2002). Misfolded proteins are competent to mediate a subset of the responses to heat shock in Saccharomyces cerevisiae. J Biol Chem 277, 44817–44825.

61. Tye, BW, and Churchman, LS (2021). Hsf1 activation by proteotoxic stress requires concurrent protein synthesis. Mol Biol Cell 32, 1800–1806.

62. Verghese, J, Abrams, J, Wang, Y, and Morano, KA (2012). Biology of the Heat Shock Response and Protein Chaperones: Budding Yeast (Saccharomyces cerevisiae) as a Model System. Microbiol Mol Biol Rev 76, 115–158.

63. Wang, J, and Sevier, CS (2016). Formation and reversibility of BiP protein cysteine oxidation facilitate cell survival during and post oxidative stress. J Biol Chem 291, 7541– 7557.

64. Wang, Y, Gibney, PA, West, JD, and Morano, KA (2012). The yeast Hsp70 Ssa1 is a sensor for activation of the heat shock response by thiol-reactive compounds. Mol Biol Cell 23, 3290–3298.

65. West, JD, Wang, Y, and Morano, KA (2012). Small molecule activators of the heat shock response: Chemical properties, molecular targets, and therapeutic promise. Chem Res Toxicol 25, 2036–2053.

66. Winterbourn, CC, and Hampton, MB (2008). Thiol chemistry and specificity in redox signaling. Free Radic Biol Med 45, 549–561.

67. Winzeler, EA et al. (1999). Functional characterization of the S. cerevisiae genome by gene deletion and parallel analysis. Science 285, 901–906.

68. Zheng, X, Krakowiak, J, Patel, N, Beyzavi, A, Ezike, J, Khalil, AS, and Pincus, D (2016). Dynamic control of Hsf1 during heat shock by a chaperone switch and phosphorylation. ELife 5.

